# Mechanism of USP21 autoinhibition and histone H2AK119 deubiquitination

**DOI:** 10.1101/2025.04.17.649347

**Authors:** Sanim Rahman, Michael T. Morgan, Chad W. Hicks, Alexander Gwizdala, Cynthia Wolberger

## Abstract

Monoubiquitinated histone H2A lysine 119 (H2AK119ub) is a signature modification associated with transcriptional silencing and heterochromatin formation. Ubiquitin-specific protease 21 (USP21), one of four major deubiquitinating enzymes (DUBs) that target H2AK119ub, plays critical roles in diverse cellular processes^1–4^. The molecular mechanisms by which USP21 specifically deubiquitinates H2AK119ub and is regulated is unknown. USP21 contains a C-terminal USP catalytic domain, preceded by an N-terminal intrinsically disordered region (IDR). We determined the cryo-EM structure of the USP21 catalytic domain bound to an H2AK119ub nucleosome, which reveals a recognition mode that differs from that of two other H2AK119-specific DUBs, Polycomb repressive complex^5^ and USP16^6^. We unexpectedly discovered that the N-terminal intrinsically disordered region (IDR) of USP21 inhibits the enzyme’s activity. Using AlphaFold-Multimer to perform a virtual screen of USP21 interactors, we identified kinases that phosphorylate the USP21 IDR and thereby relieve autoinhibition. Modeling of USP21 using AlphaFold3 suggests a structural model explaining the mechanism of autoinhibition. AlphaFold analysis of other ubiquitin-specific proteases suggests that phosphorylation-regulated autoinhibition may be a feature of multiple USP enzymes. These findings shed light on the molecular mechanisms of H2AK119 deubiquitination and reveal a novel mode of phosphorylation-dependent DUB autoregulation.

## Introduction

Reversible post-translational modification of histone proteins plays a central role in regulating eukaryotic gene transcription. Monoubiquitination of histone H2A K119 by the Polycomb repressive complex 1 (PRC1) is a defining modification associated with transcription silencing in multicellular organisms (K119 in vertebrates, K118 in *Drosophila*, K121 in *Arabidopsis*)^7,8^. In humans, deubiquitination of H2AK119 is mediated by four deubiquitinating enzymes (DUBs): BAP1^9^, MYSM1^10^, USP16^11^, and USP21^2^. Each H2AK119-specific DUB regulates distinct cellular processes and is regulated differently.

USP21 facilitates the activation of genes during transcriptional reprogramming by deubiquitinating H2AK119ub^4^. Deubiquitination of histone H2AK119ub by USP21 also promotes transcription activation through *trans-*histone crosstalk in hepatocytes, by promotion of H3K4 methylation by the MLL1 complex and concomitant activation of genes associated with hepatocyte regeneration^2^. In addition to its role in deubiquitinating H2AK119, USP21 disassembles K48-linked ubiquitin chains on AIM2^12^, BRCA2^13^, MEK2^14^, and Nanog^1^, thereby rescuing them from proteasomal degradation. USP21 also disassembles K63-linked ubiquitin chains on RIP-1 to inhibit NF-κB signaling^15^, as well as K27/K63-ubiquitin linkages on STING to downregulate innate immune response to viral DNA^16^. USP21’s activity on multiple substrates allows it to play a central role in various cellular processes, ranging from inflammasome activation to stem cell pluripotency.

Previous structural studies of USP21 focused on complexes of the C-terminal USP-family catalytic domain bound to di-ubiquitin^17^ or an inhibitor^18^, but the manner by which USP21 recognizes its ubiquitinated nucleosomal substrate was unknown. USP21 also contains a ∼200 residue N-terminal intrinsically disordered region (IDR) containing a nuclear export sequence^19,20,19,20^, but the role of the IDR, if any, in modulating USP21 enzymatic activity was unknown. We report here the structure of the USP21 catalytic domain bound to H2AK119-ubiquitinated nucleosomes. The structure reveals that USP21 simultaneously recognizes the nucleosome acidic patch and nucleosomal DNA at superhelical location (SHL) 6.5. Structural and AlphaMissense analysis of cancer mutations in USP21’s catalytic domain provides mechanistic insight into the effect of various missense mutations on USP21’s stability and deubiquitinase activity. In solution-based experiments with the intact USP21 protein, we unexpectedly discovered that the N-terminal IDR autoinhibits USP21 activity. Modeling with AlphaFold3 suggested that the IDR may occlude the ubiquitin-binding pocket of USP21, potentially inhibiting enzymatic activity. By performing an in silico screen of 152 known USP21 interactors using AlphaFold-Multimer, we identified several kinases predicted to interact with IDR residues that were known USP21 phosphorylation sites. Biochemical assays confirmed that phosphorylation of residues in the USP21 IDR by MARK1 and PRKCI relieves autoinhibition. Analysis of AlphaFold3 models of other DUBs suggest that this autoinhibition mechanism may be present in other USP family members, potentially explaining how these enzymes may be regulated.

## Results

### Structural overview of USP21 bound to a H2AK119-ubiquitinated nucleosome

In order to learn how USP21 specifically deubiquitinates H2AK119ub, we determined the cryo-EM structure of the USP21 catalytic domain (196-565) bound to a nucleosome with ubiquitin linked to H2A K119 via a non-hydrolyzable dichloroacetone (DCA) linkage^21^ (Figure 1A,B). Due to the complex’s high affinity for the carbon support, we collected two datasets on the sample prepared on graphene oxide grids to obtain a sufficient number of particles. The final map had an overall resolution of 2.98 Å, with USP21 and ubiquitin resolved at a moderate resolution of 5.33 Å (Supplementary Figures 1 and 2). Regions of the USP21 catalytic domain that were unresolved in previously reported crystal structures remained ambiguous in our maps (residues 196-202, 320-349 and 559-565)^17,18^, indicating conformational plasticity of these regions. Nonetheless, we were able to unambiguously position the crystal structures of USP21 and ubiquitin into the map and model interactions between USP21 and the nucleosome (Figure 2A,B).

**Figure 1.**
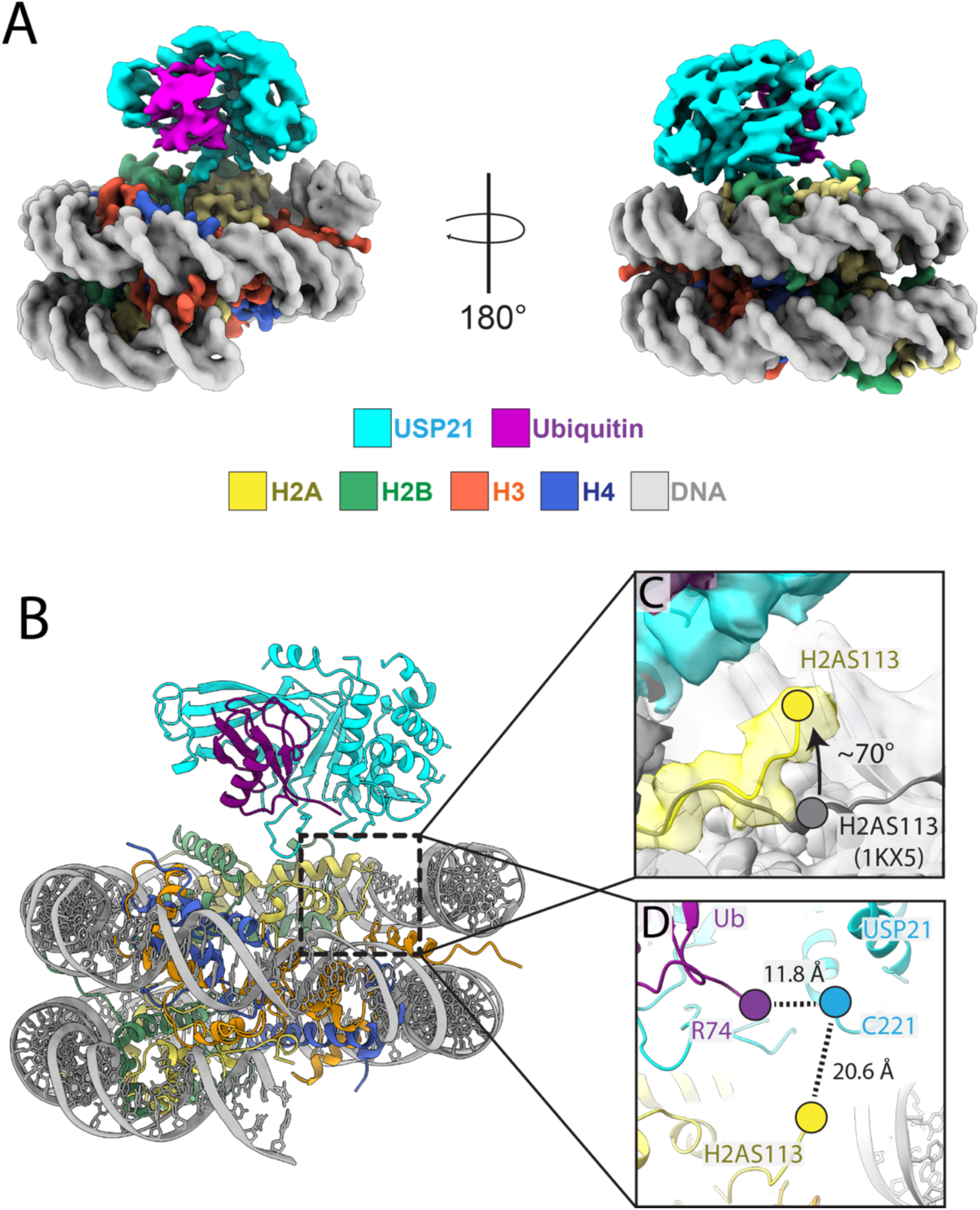
Structural overview of USP21 bound to a H2AK119-ubiquitinated nucleosome. (A) Cryo-EM reconstruction of USP21 bound to a H2AK119-ubiquitinated nucleosome (2.94 Å). (B) Cartoon representation of model. (C) Zoom on the H2A c-terminal tail in panel B. The C-terminal of H2A is lifted 70° up from the crystal structure of the nucleosome (PDB-1KX5 in gray) and positioned towards the catalytic site of USP21 (yellow). The last residue of the model (H2AS113) is shown for comparison. (D) Zoom on the USP21 catalytic site depicting the distance between Ubiquitin R74 to USP21 C221 and H2A S113 to USP21 C221.

**Figure 2.**
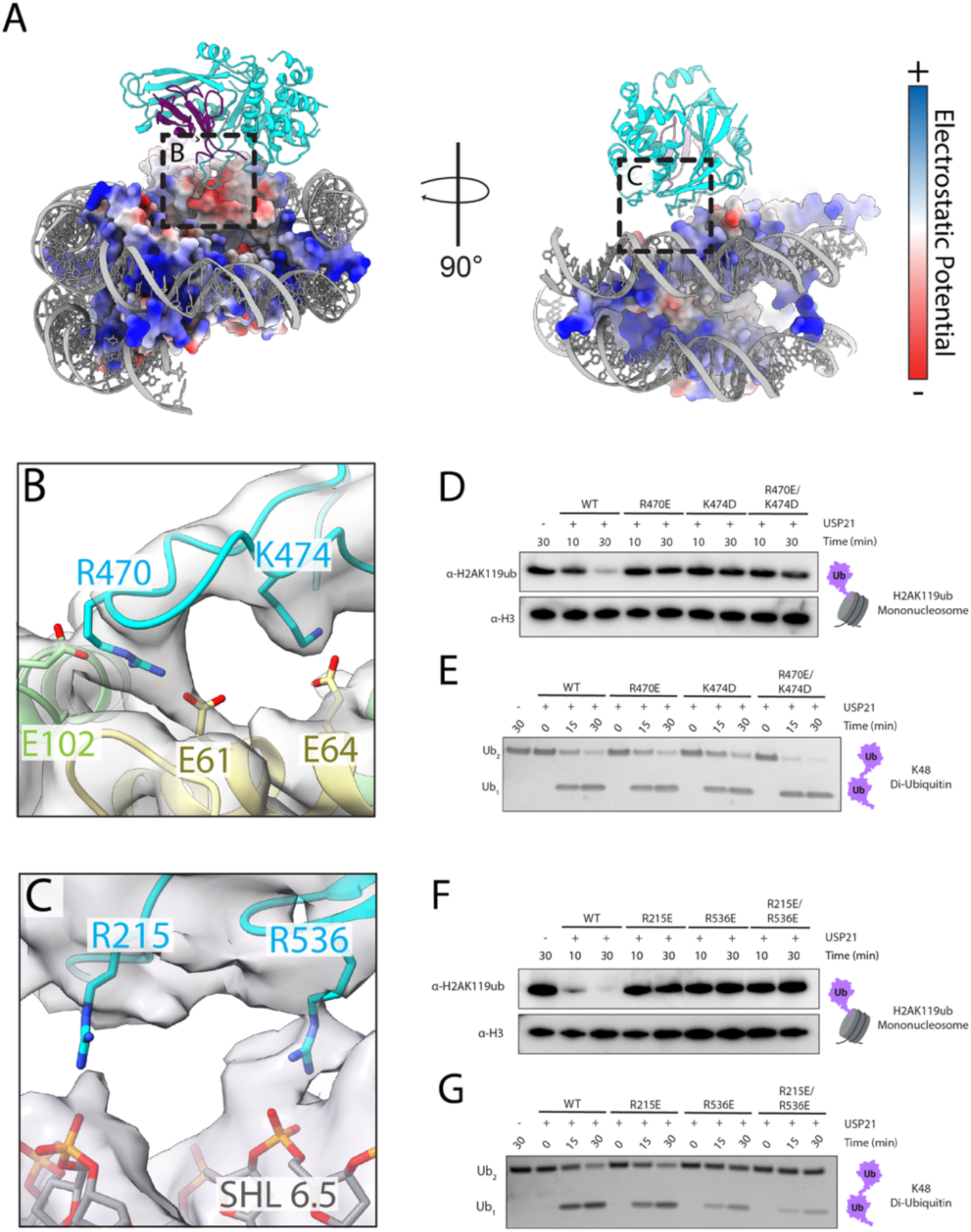
USP21’s interactions with H2AK119ub nucleosome. (A) View of USP21 with H2AK119-ubiquintated (H2AK119ub) nucleosome with the histone octamer depicted in an electrostatic potential surface representation. (B) Zoom on the interaction interface between USP21 and the acidic patch in panel A. (C) Zoom on the interaction interface between USP21 and nucleosomal DNA in panel A. (D) Effect of charge-reversal mutations of USP21’s acidic patch interactions on H2AK119ub nucleosome. Western blot analysis of USP21 activity against H2AK119ub nucleosome with USP21 WT, R470E, K474D, and R470E/K474D constructs. (E) Effect of charge-reversal mutations of USP21’s acidic patch interactions on K48-linked di-ubiquitin (K48-Ub_2_) cleavage. Coomassie blue analysis of USP21 activity against K48-Ub_2_ with USP21 WT, R470E, K474D, and R470E/K474D constructs. (F) Effect of charge-reversal mutations of USP21’s DNA interactions on H2AK119ub nucleosome. Western blot analysis of USP21 activity against H2AK119ub nucleosome with USP21 WT, R215E, R536E, and R215E/R536E constructs. (G) Effect of charge-reversal mutations of USP21’s acidic patch interactions on K48-linked di-ubiquitin (K48-Ub_2_) cleavage. Coomassie blue analysis of USP21 activity against K48-Ub_2_ with USP21 WT, R215E, R536E, and R215E/R536E constructs.

The structure shows that the H2A C-terminus adopts a conformation that enables the ubiquitin conjugated to K119 to bind USP21. Ubiquitin binds to the enzyme in a manner virtually identical to that seen in the crystal structure of USP21 bound to ubiquitin-aldehyde^17^ (Supplementary Figure 3). In addition to the density for ubiquitin, we identified a short stretch of residues at the C-terminus of histone H2A (residues 109 – S113) that is oriented towards the active site of USP21 (Figure 1C). The conformation of these H2A residues differs from that in the crystal structure of an unmodified nucleosome (PDB ID: 1KX5)^22^, which shows the C-terminal tail of histone H2A through residue K128 laying flat on the surface of the histone octamer near the nucleosome dyad (Figure 1C). Model-building shows that conformation of H2A in the unmodified nucleosome would not allow a ubiquitin linked to K119 to binds to USP21. Thus, for USP21 to engage ubiquitin that is covalently linked to K119, histone H2A must undergo a conformational rearrangement, lifting it off the histone octamer surface. This positions S113 to be 20.6 Å away from the USP21 catalytic cysteine (C221). Ubiquitin was modelled up to residue R74, placing it 11.8 Å away from C221 of USP21 (Figure 1D). The positioning of the last visible residues of histone H2A and ubiquitin are consistent with the presence of an isopeptide linkage between H2AK119 and ubiquitin within the active site of USP21. This parallels similar observations of the PR-DUB complex bound to H2AK119ub, where an even more dramatic unfolding of the H2A docking domain is required for BAP1 to engage with H2AK119ub^23^.

### USP21 interacts with the nucleosome acidic patch and DNA

USP21 is anchored to the nucleosome by a pair of basic residues, R470 and K474, that interact with the conserved nucleosome acidic patch. R470 serves as a canonical arginine anchor (Figure 2B), as has been observed for many chromatin-binding factors that engage the nucleosome acidic patch^24,25^. Specifically, R470 is poised to interact with both H2A E61 and H2B E102 of the acidic patch, as we were able to observe strong connecting density between the three residues (Figure 2B). In addition, we identified density for the K474 side chain and modeled it to interact with H2A E64, for which there is partial density for the sidechain. To test the importance of the acidic patch contacts for USP21 deubiquitination of H2AK119ub, we assayed the activity of single and double charge-reversal mutants of USP21 residues R470 and K474 on nucleosomes containing ubiquitin conjugated to H2AK119 via a native isopeptide linkage. As shown in Figure 2D, both single and double substitutions of R470 and K474 abolished the activity of USP21 on H2AK119ub. To rule out the possibility that these mutations disrupt USP21’s overall catalytic activity, we tested the effects of R470E, K474D, R470E/K474D substitutions on USP21 cleavage of K48-linked di-ubiquitin (Figure 2E). All USP21 substitution at residues implicated in binding the nucleosome acidic patch mutations had activity similar to the wild type enzyme in this assay, indicating that R470 and K474 are required for interactions with the nucleosomal substrate and not for overall enzymatic activity.

In addition to the interaction of USP21 with the nucleosome acidic patch, there is density bridging USP21 and nucleosomal DNA at SHL 6.5, suggesting that there is at least one point of contact between USP21 and the negatively charged sugar-phosphate backbone of DNA (Figure 2C). We modeled the connecting density as a contact between R536 and the DNA backbone and used model building to identify R215 as an additional potential point of contact. We tested the activity of charge-reversal mutants of R215 and R536 on H2AK119ub nucleosomes and found that both single and double substitutions at these residues abrogated USP21 activity on H2AK119ub (Figure 2F). To test whether these mutations specifically affected USP21 activity on H2AK119ub, we assayed the ability of the mutant proteins to cleave K48-linked di-ubiquitin (Figure 2G). We found that, while R215E displayed similar activity to the wild type protein, both R536E and R215E/R536E have reduced activity on di-ubiquitin, although the decrease in activity is modest as compared to the effects of these mutations on deubiquitinating H2AK119ub. Further inspection of R536 suggests that this residue may potentially be important for maintaining the structural integrity of the USP21 catalytic site, which could explain its substrate-independent effect on activity (Supplementary Figure 4).

The structure provides insights into the effects of cancer-associated mutations in USP21 that would not be predicted to simply inactivate the enzyme by destabilizing the fold of the catalytic domain. Of the 373 USP21 mutations found in the Catalogue of Somatic Mutations in Cancer (COSMIC) database and visualized in ProteinPaint^26,27^ (Supplementary Figure 5), 100 are located within the catalytic domain of USP21 (residues 212-565) (Figure 3B). Of these, Q300H, S391F, E395K, and Y519C, are predicted to disrupt polar contacts with ubiquitin (Figure 3C). Mutations at R470Q, G512F, and V514, would disrupt interactions with the nucleosome acidic patch or cause a steric clash with the H2B C-terminal helix while R215Q and R536C/L would abolish interaction of USP21 with the nucleosome DNA (Figure 3C). Interestingly, we identified E276K as a mutation that could potentially improve H2AK119ub recognition by enhancing USP21’s affinity for the nucleosome DNA.

**Figure 3.**
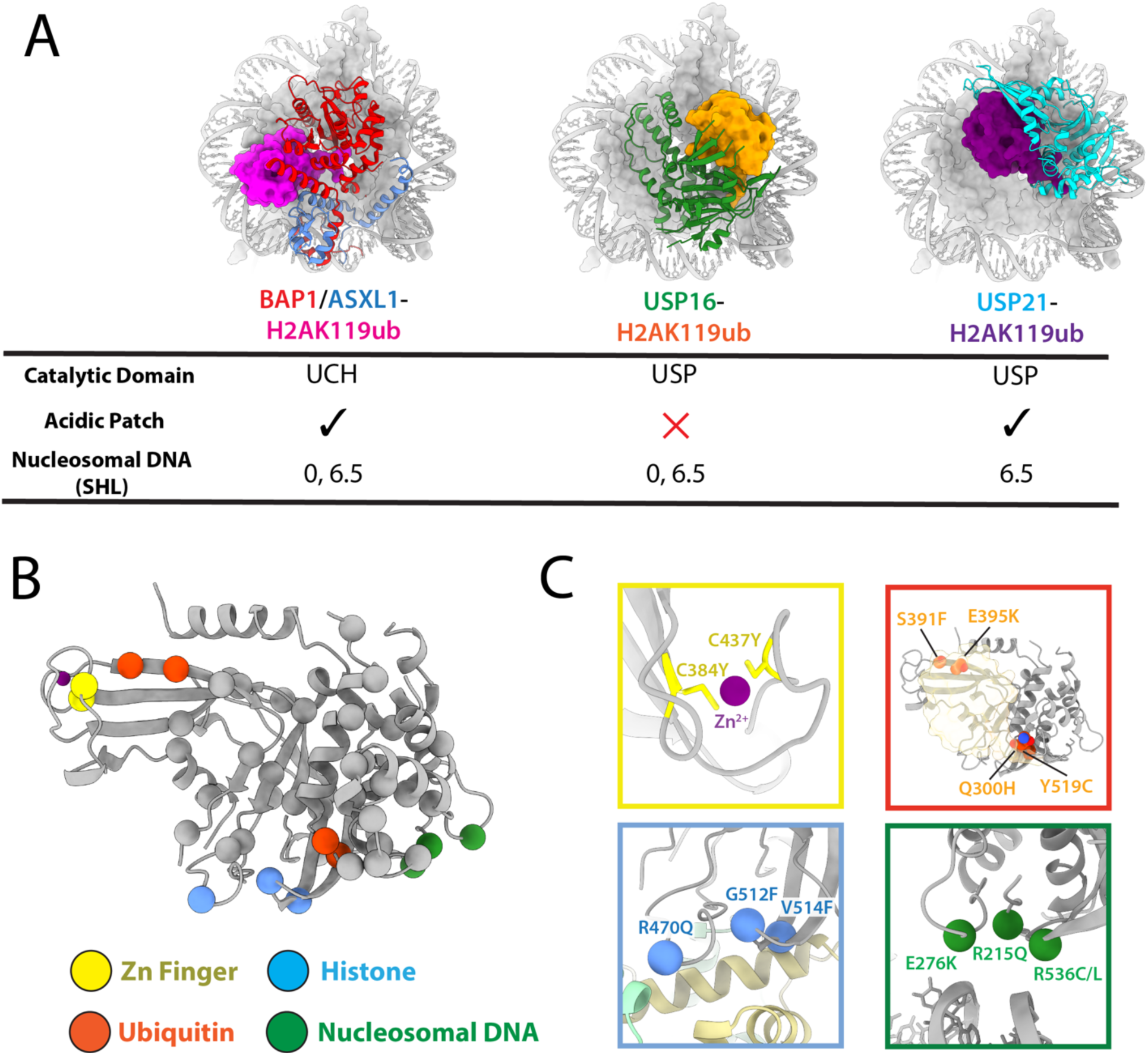
Diversity of DUB engagement of H2AK119ub and mapping of USP21 mutations. (A) Structural comparisons of H2AK119ub engagement by PR-DUB, USP16, and USP21, highlighting differences in nucleosome engagement and ubiquitin positioning. (B) Structural mapping of USP21 missense mutations. (C) Zoom-in on USP21 missense mutations and their predicted defects in USP21 activity and structure.

### Comparison of USP21 binding with other H2AK119-specific DUBs

Despite their common substrate, the three H2AK119-specific enzymes, USP21, USP16 and the PR-DUB complex^5,6,23^ have divergent modes of binding their ubiquitinated nucleosome substrate. In each structure, ubiquitin is in a different position and the enzymes are positioned in a different manner on the nucleosome (Figure 3A). The differences are particularly notable in comparing USP21 and USP16, as both enzymes contain a conserved USP catalytic domain, whereas the BAP1 catalytic domain belongs to the UCH family. Nonetheless, BAP1 and USP21 bind the nucleosome acidic patch with a canonical arginine anchor^24^, whereas USP16 does not contact the acidic patch. In addition, although all three DUBs contact the DNA at SHL 6.5, both PR-DUB and USP16 have a second contact point at SHL 0 that is absent in the USP21 complex.

To better understand the structural basis for differences between USP16 and USP21 recognition of H2AK119ub despite the similarity in their catalytic domains, we examined the potential contacts that could be formed by USP21 and USP16 in each of the complexes with H2AK119ub nucleosomes (Supplementary Figure 6). Whereas USP16 utilizes basic residues to interact with DNA at SHL 0 and 6.5 as well as with the C-terminal helix of histone H2B, USP21 does not contain any basic residues at these positions. In addition, USP16 contains a unique structured insertion that mediates binding to the C-terminal helix H2B that is absent in USP21 (Supplementary Figure 6A,B). Conversely, USP16 lacks key basic residues that position USP21 on H2AK119ub nucleosomes, including the arginine anchor and arginine residues that contact DNA in the USP21 complex (Supplementary Figure 6C,D). These differences in nucleosome engagement highlight the importance of both DUB structure and sequence in substrate recognition.

### USP21 is autoinhibited by its N-terminal disordered region

In preparing USP21 protein for structural studies, we expressed and purified full-length USP21 (residues 1-565), as well as a series of fragments containing successive N-terminal truncations of the IDR (Figure 4A). Activity assays of the minimal catalytic domain (196–565) using the ubiquitin-aminomethylcoumarin (Ub-AMC) fluorescent reporter showed robust cleavage activity, as has been previously reported^17^. However, we unexpectedly found that the full-length enzyme exhibited essentially no activity in these assays (Figure 4B). N-terminal truncation mutants lacking residues 1-47 or 1-121 of the IDR similarly had little to no deubiquitinating activity, whereas further truncations removing residues 1-158 and 1-184 of the IDR had 53% and 77% the activity of the USP21 catalytic domain, respectively (Figure 4B). To verify that the observed autoinhibition was not specific to the Ub-AMC assay, we tested the ability of the USP21 constructs to cleave K48-linked di-ubiquitin chains, which had been previously reported as a USP21 substrate^1,12–14^. As shown in Figure 4C, the full-length protein had no deubiquitinating activity in this assay, whereas successive deletions showed increasing activity, with maximum activity observed for the minimal catalytic domain. We also compared the activity of full-length USP21 on nucleosomes containing H2AK119ub with that of the minimal catalytic domain. The full-length protein showed only minimal activity over a 1-hr time course, while the catalytic domain alone fully deubiquitinated nucleosomal histone H2A over the same period (Fig. 4D). Intact USP21 containing the full-length IDR also showed no detectable binding of USP21 to H2AK119-ubiquitinated nucleosomes (Figure 4E), whereas the catalytic domain alone bound to ubiquitinated nucleosomes with a K_d_ of around 0.6 μM. Taken together, these experiments indicate that the USP21 N-terminal IDR inhibits the enzyme’s deubiquitinating activity in a substrate-independent manner.

**Figure 4.**
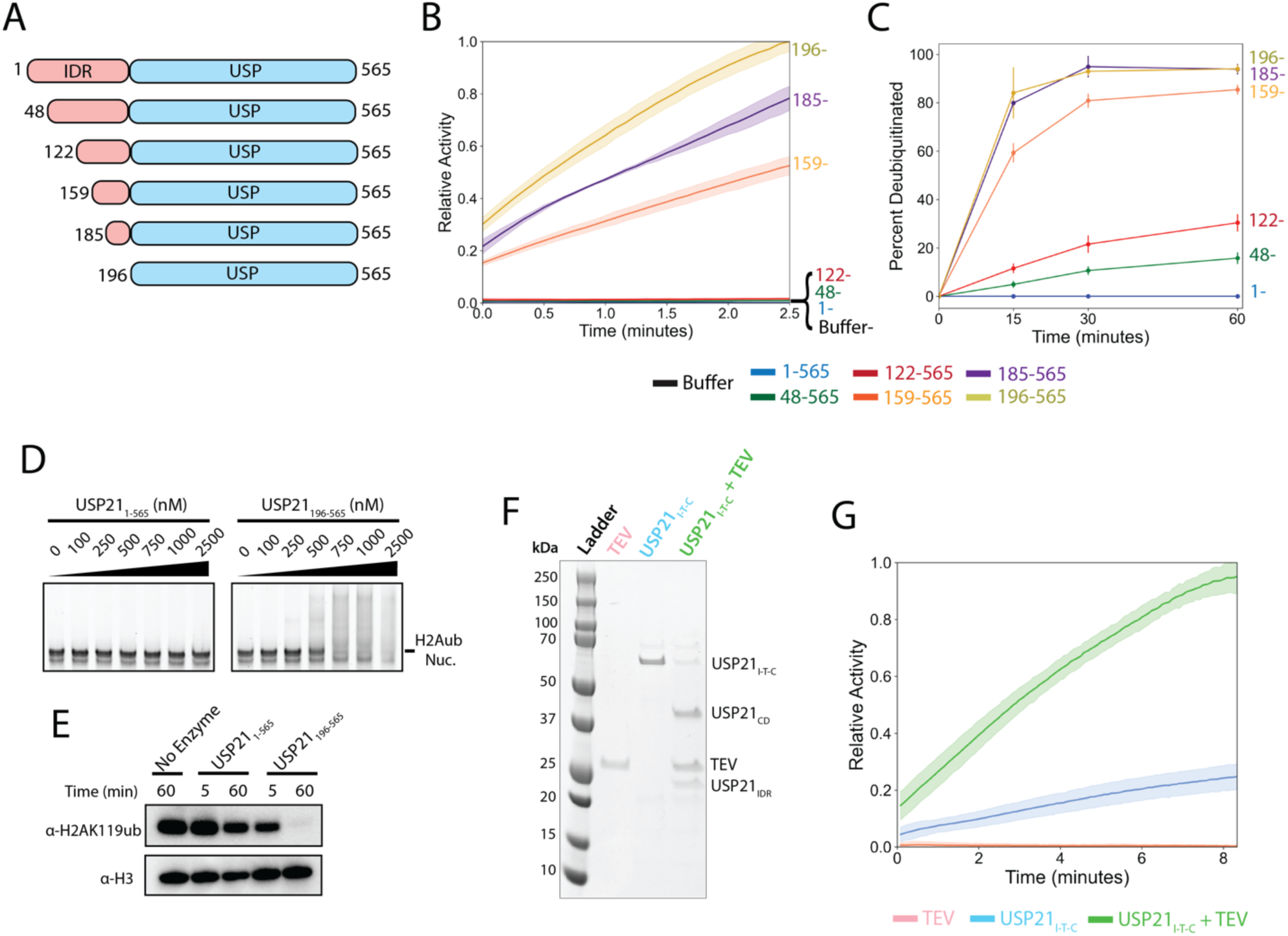
The USP21 IDR is an autoinhibitory module. (A) Schematic of USP21 and truncations used in this study. Truncations were made within the disordered N-terminal region, upstream from the catalytic domain. (B) Time-course curve of Ub-AMC cleavage by full-length USP21 and N-terminal truncations. (C) Effect of N-terminal truncations on USP21’s activity against K48-linked di-ubiquitin. (D) Electrophoretic mobility shift assay of full-length USP21 and catalytic domain to H2AK119ub nucleosome. (E) Western blot analysis of H2AK119-deubiquitination activity by full-length USP21 and catalytic domain. (F) Design of TEV-cleavable USP21 construct (USP21_I-T-C_) and Coomassie blue analysis of USP21_I-T-C_ cleavage upon addition of TEV protease. (G) Time-course curve of Ub-AMC cleavage by TEV protease, USP21_I-T-C_, and USP21_I-T-C_ pre-incubated with TEV protease.

Although the full-length USP21 protein appeared to be well-behaved in solution, we wished to rule out the possibility that the full-length protein was inactive due to misfolding. We therefore engineered a full-length USP21 construct in which residues 190-195 were substituted to create a Tobacco Etch Virus (TEV) protease cleavage site (USP21_I-T-C_). This design made it possible to assay the activity of the same USP21 protein before and after cleavage with TEV protease to separate the IDR from the catalytic domain. As shown in Figures 4F and 4G, preincubating USP21_I-T-C_ with TEV protease led to a dramatic increase in activity as assayed by cleavage of Ub-AMC, indicated that the presence of the IDR inhibits USP21 activity (Figure 4G).

Interestingly, the USP21_I-T-C_ protein exhibits somewhat higher activity than the native full-length protein, suggesting that the altered amino acid sequence in residues 190-195, which are immediately N—terminal to the catalytic domain, may in some way reduce the ability of the IDR to autoinhibit USP21.

### In silico pulldown identifies kinases and phospho-binding proteins as potential IDR interactors

Since USP21 must be active in order to carry out its functions, we considered several possible mechanisms by which autoinhibition could be relieved. As has been found for other DUBs, autoinhibition can be regulated by oligomerization state^28–30^, phosphorylation^31,32^, or interaction with partner proteins^33,34^. Since full-length USP21 migrates as a monomer in size-exclusion chromatography (Supplementary Figure 7), we therefore speculated that USP21 may be regulated by either phosphorylation or by binding to partner proteins.

To identify potential binding partners or enzymes that interact with the USP21 IDR, we used an in-silico approach to rapidly screen for proteins that could potentially relieve autoinhibition of USP21. A total of 152 interacting partners of USP21 have been identified across multiple studies ^35–39^ and are tabulated in the BioGRID database^40^. Using AlphaFold-Multimer^41^ as implemented in the AlphaPulldown^42^ pipeline, we predicted structures for all pairwise interactions between USP21 and its interactors (Figure 5A). To identify proteins that interacted with the USP21 IDR, we carried out pairwise predictions for all 152 interactors with the full-length USP21 protein (residues 1-565) and with the catalytic domain alone (residues 196-565). We rationalized that interactors that engage USP21 IDR would have a significantly higher interface predicted

**Figure 5.**
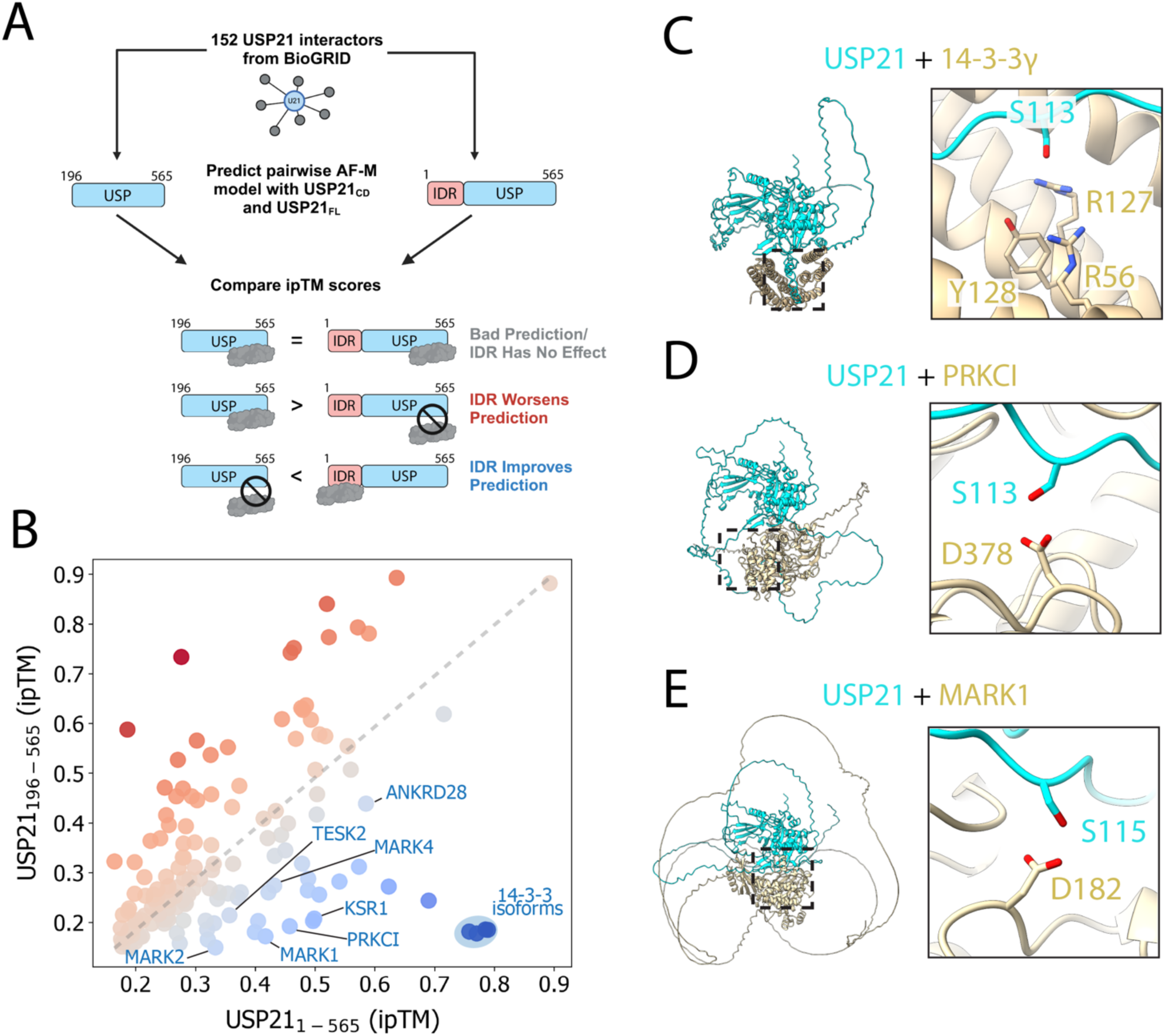
In silico screening of USP21_IDR_ interactors with AlphaFold-Multimer. (A) Schematic describing pipeline of screening and identifying interactors of USP21_IDR_ using AlphaFold-Multimer. (B) Comparison of ipTM score of all interactors predicted with either full-length USP21 or the catalytic domain. (C-E) AlphaFold-Multimer prediction of full-length USP21 with 14-3-3γ, PRKCI, and MARK1 with a zoom-in on 14-3-3γ’s phosphoserine binding pocket poised to bind to USP21_S113_ or the catalytic residues of PRKCI and MARK1 poised to phosphorylate USP21_S113_ or USP21_S115_.

Template Modeling (ipTM) score for the USP21_1-565_ model as compared to USP21_196-565_. We generated five models for each interaction, yielding over 1500 models, and used the ipTM score reported for the highest confidence model. To identify proteins predicted to interact with the USP21 IDR, we computed a ΔipTM score, which we define as the difference in ipTM between USP21_1-565_ and USP21_196-565_ for a given interactor (Supplementary Table 3). Interactors with ipTM scores of less than 0.35 for both USP21 constructs were classified as low-confidence due to the inability of AlphaFold to predict the interaction. This comprised 75 of the 152 interactors. An additional 57 interactors showed either no significant difference (-0.15<ΔipTM<0.15), indicating that the IDR was not important for interactions with USP21, or had a significant negative ΔipTM score (ΔipTM<-0.15), indicating that the presence of the IDR worsened the prediction. The remaining 20 interactors had an improved prediction with the IDR, which we considered as a ΔipTM>0.15, and were considered candidate USP21 IDR interactors.

To validate the utility of using ΔipTM as a metric for identifying proteins that bind to the USP21 IDR, we assessed whether known interactors were identified in our in silico screen. Previous work had shown that the nuclear export sequence of USP21 lies within the IDR (residues 134-152). We found that Exportin was one of the 20 interactors and was predicted to form a canonical interaction with the nuclear export sequence (LXXXLXXLXL) with high confidence (Supplementary Figure 8C,D). Another USP21 interactor identified in our screen was the MARK2 kinase (Supplementary Figure 8E,F), which had previously been shown to interact with IDR residues 48-121^36^. Overall, about half the proteins predicted to interact with the USP21 IDR were either kinases, phosphatases, or 14-3-3 proteins,which bind phosphoserine and phosphothreonine^43^, suggesting that the IDR may be regulated by phosphorylation (Figure 5B-E and Supplementary Figure 8). Consistent with this hypothesis, the predicted complexes with the 14-3-3 isoforms showed S113 of USP21 bound to the canonical phosphoserine binding pocket of 14-3-3 proteins (Figure 5C and Supplementary Figure 8A,B). An analysis of potential binding motifs with the 14-3-3 Pred server^44^ showed USP21 residue S113 to be one of five predicted high confidence 14-3-3 binding sites (Supplementary Figure 9A,B). Our predictions highlighted multiple kinases (MARK1, MARK2, MARK4, and PRKCI) poised to phosphorylate USP21 at S113 and S115 (Figure 5D,E and Supplementary Figure 8E,F), both of which had been shown to be heavily phosphorylated in both HEK293T (91.5% and 97.3%, respectively) and HeLa cells (57.9% and 82.5%, respectively)^36^. The residues flanking S113 and S115 were analyzed using Kinase Library^45,46^ to identify high-confidence phosphorylation sites for specific USP21 IDR interactors. For PRKCI (PKCI), S113 was ranked at the 98.4^th^ percentile of substrates (Supplementary Figure 9C,D). For MARK1, S115 was ranked at the 98.5% percentile and was the top-ranked kinase for the motif (Supplementary Figure 9E,F). Taken together, our results suggest that autoinhibition of USP21 may be regulated by phosphorylation-dependent mechanisms.

The IDR must in some way interfere with USP21 substrate binding or catalysis in order to inhibit deubiquitinating activity. To identify a potential self-interaction between the IDR and the catalytic domain, we predicted the structure of full-length of USP21 using AlphaFold3 (Figure 6A)^47^. Based on the Predicted Aligned Error (PAE), we identified a high-confidence interaction involving residues 108-117 with the catalytic domain of USP21 (Figure 6B,C). This high confidence interaction positions the IDR in the ubiquitin binding site in the USP21 catalytic domain. Notably, residues S113 and S115, which are known phosphorylation sites, are involved in the interaction. Based on this model, we hypothesized that the IDR autoinhibits USP21 by blocking access the ubiquitin binding interface and that this interaction could be disrupted by phosphorylation. We tested this hypothesis by using AlphaFold3 to predict the structure of USP21 with S113 and S115 phosphorylated, as well as with a number of other known phosphorylation sites, to see if phosphorylation of any residues resulted in a model in which the IDR no longer occluded the ubiquitin-binding site (Figure 6D,E and Supplementary Figure 10)^36^. The only predictions that yielded a putative uninhibited model contained phosphorylated S93, S113, and S115. Notably, these are residues previously identified as highly phosphorylated in HeLa and HEK293 cells^36^.

**Figure 6.**
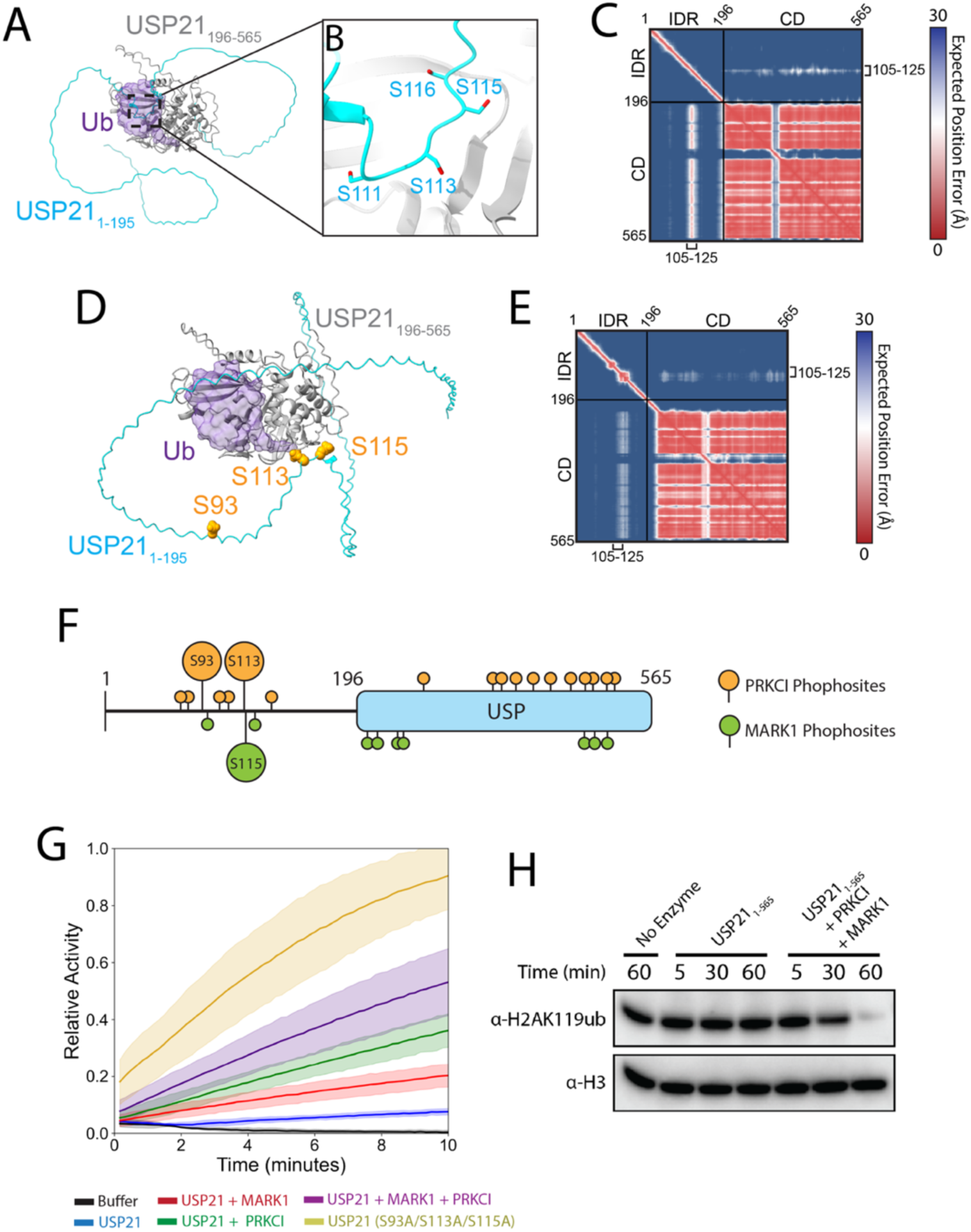
AlphaFold3-guided biochemistry reveals mechanism of autoinhibition relief by phosphorylation. (A) AlphaFold3 structure prediction of full-length USP21 (apo) shown in respect to ubiquitin binding (purple). (B) S111, S113, S115, and S116 positioned at the ubiquitin-binding binding interface of USP21’s catalytic domain. (C) Predicted Aligned Error (PAE) of full-length USP21 (apo), highlighting the high-confidence spatial positioning of USP21_105-125_ with respect to the catalytic domain. (D) AlphaFold3 structure of full-length USP21 phosphorylated at S93, S113, and S115 (apo) shown with respect to ubiquitin binding. (E) PAE of full-length USP21 phosphorylated at S93, S113, and S115 (apo), highlighting the low confidence spatial positioning of USP21_105-125_ which is also positioned away from the ubiquitin binding interface. (F) Phosphorylated residues of USP21_IDR_ upon treatment of PRKCI or MARK1, identified by mass spectrometry. (G) Time-course curve of Ub-AMC cleavage by full-length USP21 when treated with PRKCI and/or MARK1 and USP21_S93A/S113A/S115A_. (H) Western blot analysis of H2AK119-deubiquitination activity by full-length USP21 with and without treatment of PRKCI and MARK1.

### Phosphorylation by PRKC1 and MARK1 relieves USP21 autoinhibition

Our in silico modeling suggested that phosphorylation of residues S93, S113 and S115 in the USP21 IDR could relieve autoinhibition of the enzyme and identified PKC1 and MRK1 as the putative kinases. To confirm that S93, S113 and S115 were key residues mediating autoinhibition, we mutated S93, S113 and S115 to alanine (USP21-3A) and found that the mutant enzyme was highly active in cleaving Ub-AMC (Figure 6G). To test the prediction that these residues could be phosphorylated by PRKC1 and MARK1, we incubated full-length USP21 with these enzymes and identified phosphorylation sites using mass spectrometry (Figure 6F and Supplementary Table 4). We found that PRKCI was able to phosphorylate S113 while MARK1 phosphorylated S115, validating the predictions from AlphaFold-Multimer. In addition, PRKCI phosphorylated S93, suggesting that treating USP21 with both kinases could fully activate the protein. To test whether phosphorylation relieved autoinhibition, we measured the activity of USP21 after treatment with PRKCI, MARK1 or both (Figure 6G). Incubation with each kinase increased USP21 cleavage activity on Ub-AMC, while treating USP21 with both kinases resulted in the highest activity. To rule out substrate-dependent effects, we tested the activity of phosphorylated USP21 on H2AK119ub on nucleosomes and observed a similar increase enzymatic activity on this substrate (Figure 6H). To confirm that the activity increase was due to USP21 phosphorylation and not from any DUB contaminants present with PRKCI or MARK1, we incubated each kinase with Ub-AMC and observed no cleavage activity (Supplementary Figure 11). Together, these findings support the model that S93, S113 and S115 are critical residues regulating USP21 autoinhibition and that phosphorylation of these residues activates the enzyme’s activity.

## Discussion

Our structural study reveals unanticipated plasticity in the recognition of nucleosomes by enzymes that remove ubiquitin from histone H2AK119. Although USP21 shares a common substrate with the previously studied enzymes, USP16 and PR-DUB, it differs from both in its interaction with nucleosomes and recognition of H2AK119ub. The finding that USP16^6^ and the PR-DUB complex^5,23^ differed from one another in the manner in which they bind to nucleosomes and recognize ubiquitin (Figure 3A) was to be expected, since USP16 belongs to the USP family of DUBs and PR-DUB belongs to the structurally distinct UCH family^48^. Moreover, USP16 is a monomeric enzyme, whereas PR-DUB is a heterodimer comprising the BAP1 catalytic subunit and ASXL1, both of which mediate contacts with the nucleosome and with ubiquitin^5,23^. Although USP21 is, like USP16, a monomeric enzyme and a member of the USP family, the mechanism by which the two enzymes specifically recognize H2AK119ub is completely different. Apart from the conserved manner in which ubiquitin binds to the USP domain, the catalytic domains are docked on the nucleosome in entirely different orientations (Figure 3A). Both catalytic domains contact the DNA but do so with different regions of the USP domain (Supplementary Figure 6). Whereas USP21 contacts the nucleosome acidic patch, a hotspot for nucleosome interactions^24^, USP16 does not. We note that the location of the K119 ubiquitination site in a conformationally flexible region of histone H2A makes it possible for proteins to bind the nucleosome in a range of orientations while simultaneously binding to the ubiquitin (Supplementary Figure 12).

Our finding that USP21 is autoinhibited by an N-terminal IDR, and that this inhibition can be relieved by phosphorylation, highlights the power of combining curated experimental databases to uncover mechanisms of enzyme regulation. By using AlphaPulldown^42^ to model complexes of experimentally identified USP21 interactors bound to the full-length enzyme as well as to a truncated version lacking the IDR, we were able to identify proteins predicted in interact with the IDR. This approach allowed us to identify phosphorylation as a likely mechanism for regulating autoinhibition by the IDR. The ability to include phosphorylated residues in modeling as implemented in AlphaFold3^47^ made it possible to generate more specific models for how phosphorylation of particular USP21 residues could activate the enzyme. Our approach to identifying IDR interactors can be applied to other systems to search for enzymes or binding partners that interact with specific protein sequences or domains.

Autoregulation of DUBs by their IDRs may be a mechanism common to multiple members of the USP family. There are a number of DUBs in this structural class containing IDRs that are heavily phosphorylated and, in some cases, are known to be activated by phosphorylation^31,32^. A visual inspection of the 51 USP DUBs in the AlphaFold2 database revealed 12 USP DUBS with IDRs predicted to contact the catalytic domain (Figure 7 and Supplementary Table 5). Models for the candidate DUBs were predicted again using AF3 to generate five models of the DUB alone and with ubiquitin bound. We extracted high PAE contacts between the IDRs and catalytic domain in the apo structure, which we defined as contacts within 5 Å with a PAE cutoff of 20 Å found in at least two of the five models. All interactions that met this threshold were superimposed on the ubiquitin-bound structure to identify contiguous regions that would compete with ubiquitin binding and then these regions were analyzed to identify known phosphorylation sites. In total, twelve DUBs containing potential autoinhibitory regions were identified using this approach (Figure 7B), including USP10 and USP37 (Supplementary Figure 13), which have been shown to be activated by phosphorylation of the IDR^31,32^. Since many classes of human enzymes contain IDRs, combining protein structure prediction tools with experimental data provides a rapid route to generating testable hypotheses regarding enzyme regulation.

**Figure 7.**
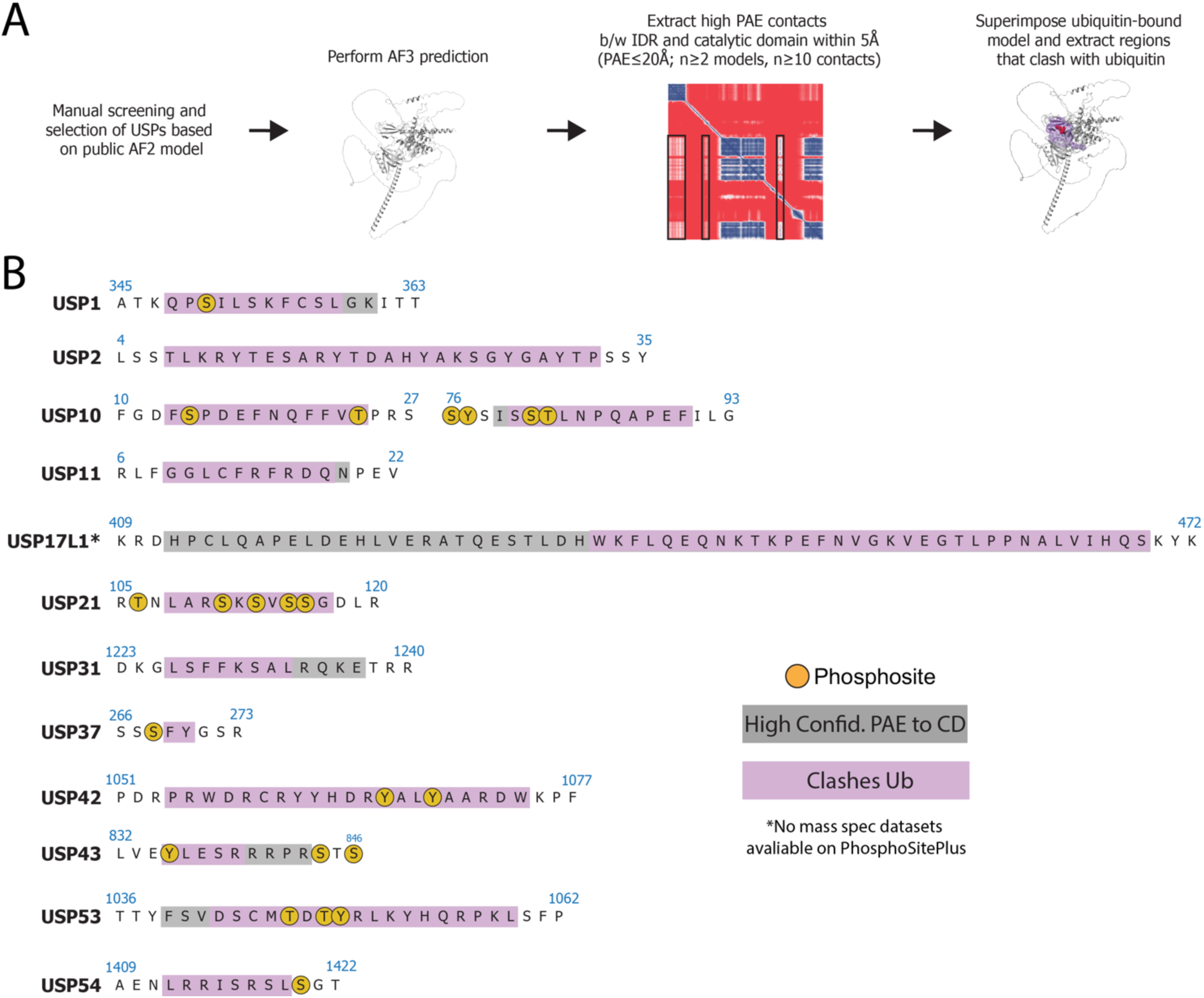
Structural informatics search for other autoinhibited ubiquitin specific protease (USP) deubiquitinases (DUBs). (A) Workflow to identify USP DUBs that contain IDRs that occlude ubiquitin binding. (B) List of DUBs and their IDR segments that are predicted to be in high spatial confidence to the ubiquitin binding interface (gray) and clash with ubiquitin binding (purple). Residues that have been identified to be phosphorylated are marked in gold.

## Methods

### Expression and purification of ubiquitin

Ubiquitin expression and purification were performed as described previously ^49^. Briefly, His-TEV-Ubiquitin containing a G76C substitution was expressed in Rosetta2 cells. Ubiquitin was first purified by Ni-NTA affinity chromatography and then dialyzed into ammonium acetate pH 4.5 overnight. Dialyzed protein was loaded onto a SP-HP cation exchange column (GE), equilibrated in ammonium acetate pH 4.5, and eluted over a linear gradient to 750 mM NaCl. Fractions were dialyzed in ubiquitin storage buffer (10 mM Tris-HCl pH 7.8, 1 mM TCEP), flash frozen, and stored at -80°C.

### Expression and purification of histones

Histone purifications were performed as described previously^50,51^. *Xenopus laevis* histone plasmids containing H2A, H2B, H3 and H4 were a gift from Greg Bowman.

Histones were expressed in Rosetta2-pLysS cells and grown in 2X YT media. Cells were induced with 1 mM IPTG at an OD600 between 0.4-0.6 and further grown for 2-4 hours at 37°C before harvested by centrifugation. Pellets were resuspended in Histone Wash Buffer (50 mM Tris pH 7.5, 100 mM NaCl, 1 mM EDTA, 5 mM BME, 1 mM PMSF), flash frozen in liquid nitrogen, and stored at -80°C.

Resuspended pellets were thawed and lysed three times through a microfluidizer. The lysate was centrifuged and pellets were resuspended with Triton Wash Buffer (50 mM Tris pH 7.5, 100 mM NaCl, 1 mM EDTA, 5 mM BME, 1% Triton X-100). The pellet was centrifuged, supernatant removed, and resuspended in Histone Wash Buffer. Resuspended pellets were then centrifuged and had the supernatant removed before storing at -20°C. Frozen pellets were later thawed and resuspended in DMSO, followed by gradual titration of Histone Unfolding Buffer (20 mM HEPES pH 7.5, 7 M Guanidine HCl, 10 mM DTT) at room temp over one hour. Once resuspended, the lysate was centrifuged, and the supernatant was harvested. The supernatant was filtered before purified over a Sephacryl 26/60 S200 sizing column equilibrated in Histone Sizing Buffer (10 mM Tris pH 8.0, 7 M urea, 1 mM EDTA, 5 mM BME). Fractions were pooled and injected onto a tandem-column series of a HiTrap Q-XL column connected upstream of a HiTrap SP-XL column using Histone Ion Exchange Buffer A (20 mM Tris pH 7.8, 7 M urea, 1 mM EDTA, 5 mM BME). The Q-XL column was removed from the system, and then eluted over a gradient with Histone Ion Exchange Buffer B (20 mM Tris pH 7.8, 7 M urea, 1 M NaCl, 1 mM EDTA, 5 mM BME). Purified histones were dialyzed in 5 mM BME, concentrated, lyophilized, and stored at -20°C.

### Preparation of nonhydrolyzable linkage H2AK119ub

H2AK119ub containing a nonhydrolyzable dichloroacetone (DCA) linkage between H2AK119C and G76C was prepared as described^49,52^. Briefly, 100 μM of His-TEV-Ubiquitin (G76C) and equimolar H2AK119C was mixed in crosslinking buffer (50 mM Borate pH 8.1, 1 mM TCEP) and incubated at 50°C for one hour. The mixture was then cooled on ice for 10 minutes before initiating crosslinking. The reaction was initiated by the addition of 100 μM DCA and the reaction was allowed to proceed for one hour. The reaction was quenched with 50 mM BME, dialyzed into water, and then lyophilized. The lyophilized product was resuspended in Ni-NTA-6MU buffer (6M urea, 50 mM Tris pH 7.8, 500 mM NaCl, 10 mM BME, 20 mM Imidazole, 0.1 mM PMSF) and incubated with Ni-NTA resin. Unreacted H2A and crosslinked H2A-were removed in the flow through. The desired product, unreacted ubiquitin and crosslinked ubiquitin-ubiquitin was eluted in NiNTA-6MU buffer supplemented with 250 mM imidazole and dialyzed into TEV cleavage buffer (50 mM Tris-HCl pH 8.0, 0.5 mM EDTA, and 1 mM DTT). The His-tag on ubiquitin was removed by the addition of TEV protease. The product was dialyzed into water, lyophilized, and re-dissolved in NiNTA-6MU buffer. The product was passed through Ni-NTA to remove uncleaved protein and His-tagged-TEV. The flowthrough was dialyzed into water and lyophilized again. Finally, the dry protein was dissolved in 7M Guanidine-HCl and separated on a Proto 300 C4 column (Higgins Analytical) in 0.1% TFA and a gradient of 0.1%TFA + 90% acetonitrile. Fractions corresponding to pure H2AK119ub were pooled and lyophilized.

### Purification of Widom 601 DNA

The 147 bp Widom 601 DNA sequence ^53^ was produced from the pST55-16x601plasmid ^54^. The pST55-16x601 plasmid was purified and the 601 sequence was isolated as described previously^55^.

### Reconstitution of H2AK119-ubiquitinated nucleosomes

H2AK119-ubiquitinated nucleosomes were prepared essentially as described previously^52^. In brief, lyophilized histone proteins (H2AK119ub, H2B, H3, H4) were resuspended in histone unfolding buffer (20 mM Tris pH 7.5, 7 M guanidine HCl, 10 mm dithiothreitol (DTT)) and combined in an H2AK119ub:H2B:H3:H4 equimolar ratios. Combined histones were dialyzed into histone refolding Buffer (10 mM Tris pH 7.5, 2 M NaCl, 1 mM EDTA, 5 mM BME) to refold the histones into the histone octamer. The octamer was then purified using a HiLoad 16/600 Superdex 200 pg size exclusion column in histone refolding buffer.

For the reconstitution, H2AK119ub octamer, dimer, and Widom 601 DNA was combined in an octamer:dimer:DNA molar ratio of 1.1:0.2:1 in high salt buffer (10 mM Tris pH 7.5, 2 M KCl, 1 mM EDTA, 1 mM DTT) such that the final DNA concentration was 6 µM. The salt in the mixture was gradually reduced over 24 hours by salt gradient dialysis into Low Salt Buffer (10 mM Tris pH 7.5, 0.25 mM KCl, 1 mM EDTA, 1 mM DTT) to assemble nucleosome. Any precipitate was removed by centrifugation and the purity of the nucleosome sample was assessed using electrophoretic mobility shift assay (EMSA). If the nucleosome sample showed excess free DNA or higher-order species, it was additionally purified by high-performance liquid chromatography using a SK DEAE-5PW column (TOSOH biosciences) with Buffer A (10 mM Tris pH 7.5, 0.25 M KCl, 0.5 mM EDTA, 1 mM DTT) and a gradient elution of Buffer B (10 mM Tris pH 7.5, 0.6 M KCl, 0.5 mM EDTA, 1 mM DTT) using an Agilent HPLC instrument. Purified nucleosome was dialyzed into Nuc Freezing Buffer (20 mM HEPES pH 7.5, 25 mM KCl, 1 mM EDTA, 1 mM DTT, 20% glycerol), flash frozen in liquid nitrogen, and stored at - 80°C.

### Expression and purification of USP21 constructs

All USP21 constructs were expressed in BL21(DE3)Rosetta2-pLysS *E. coli* cells in 2XYT media and contains an N-terminal 6x-His-SUMO tag. The cells were induced overnight with 0.5 mM IPTG at an OD600 between 0.6-0.7 at 18-20°C. Pellets were resuspended in USP21 Lysis Buffer (50 mM Tris-HCl pH 7.5, 500 mM NaCl, 10µM of ZnCl_2_, 40 mM Imidazole, 10% glycerol, 2 mM BME) supplemented with 1mM PMSF and flash frozen in liquid nitrogen, and stored at - 80°C.

Pellets were thawed in a water bath and lysed by sonication for 2.5 minutes (5 sec on, 5 sec off) at 50% power or passed through a microfluidizer three times. Following, the lysates were then clarified by centrifugation (11,000 RPM at 4°C for 30 min). The supernatant was filtered and incubated with 3-5 mL of Ni-NTA resin for one hour at 4°C. Following, the supernatant-bead slurry was loaded over a gravity column to remove unbound protein and the beads were resuspended in Wash Buffer. Beads bound with lUSP21 constructs containing parts of the IDR (1-, 48-, 122-, 159-, 185-565) were first washed with USP21 Lysis Buffer supplemented with 20mM MgCl_2_ and 5mM ATP to remove any chaperone contaminants. For all constructs, the beads were washed twice with USP21 Lysis Buffer, in which the beads were incubated with 10 column volumes of wash buffer for 30 minutes at 4°C. Following, the beads were then resuspended in 10 mL of Elution Buffer (50 mM Tris-HCl pH 7.5, 500 mM NaCl, 10µM of ZnCl_2_, 300 mM Imidazole, 10% glycerol, 2 mM BME) for 1 hour at 4°C and eluted. Another elution was performed with only a 30-minute incubation.

Eluted protein was pooled and mixed with 1 mg of 6x-His-SENP protease to cleave off the SUMO-tag overnight at 4°C. The next day, the eluted protein was dialyzed for 4 hours at 4°C in USP21 IEX Buffer A (20 mM Tris-HCl pH 7.5, 100 mM NaCl, 10µM of ZnCl_2,_ 2 mM DTT, 10% glycerol). The sample was then loaded over a HiTrap SP-XL column with a gradient elution using USP21 IEX Buffer A and USP21 IEX Buffer B (20 mM Tris-HCl pH 7.5, 1M NaCl, 10µM of ZnCl_2,_ 2 mM DTT, 10% glycerol). Fractions of interest was concentrated and injected over a HiLoad 16/60 Superdex 75 column or Superdex 75 Increase 10/300 GL equilibrated in 20 mM Tris-HCl pH 7.5, 250 mM NaCl, 5 mM DTT, 10% glycerol. Fractions of interest were pooled together, concentrated, flash frozen in liquid nitrogen as single-use aliquots, and stored at -80°C.

### Cryo-EM sample preparation of USP21-H2AK119ub nucleosome

For assembly of the USP21-H2AK119ub complex, a final concentration of 240 nM of H2AK119ub nucleosome was incubated with 1.2 uM of USP21_196-565_ in 4.5mL of crosslinking buffer (20 mM HEPES pH 7.5, 50 mM NaCl, 1 mM DTT, 2.5mM MgCl_2_) for one hour on ice. Following, 900 uL of 0.6% glutaraldehyde crosslinker, diluted in crosslinking buffer, was added to reaction to obtain a final concentration of 200 nM H2AK119ub nucleosome, 1 uM USP21_196-565_, and 0.1% glutaraldehyde. The crosslinking reaction was allowed to continue on ice for 15 minutes before being quenched with 600 uL of 1M Tris-HCl (pH 7.5), reaching a final Tris-HCl concentration of 100 mM. The sample was then concentrated and dialyzed against sample buffer (10 mM HEPES pH 7.5, 50 mM KCl, 1 mM DTT, 2.5mM MgCl_2_) for two hours. The dialyzed sample was further concentrated before freezing. Within 30 minutes of freezing, 0.25% octyl-glucopyranoside prepared in sample buffer was added to the sample, reaching a final concentration of 0.05% of detergent.

Sample was vitrified on Quantifoil 400 mesh R1.2/1.3 containing a graphene oxide support layer prepared by the manufacturer. A 3.5 uL of USP21-H2AK119ub nucleosome sample at a concentration of 0.6 mg/mL was applied onto the grid and plunge froze at 4°C and 100% humidity using a FEI Vitrobot Mark IV.

### Cryo-EM data collection and processing

Two cryo-EM data sets were collected at the Beckman Center for Cryo-EM at the Johns Hopkins University School of Medicine, using a Titan Krios at 300 kV equipped with a Falcon 4 Direct Electron Detector with Selectris Energy Filter. The first dataset contained 10,737 micrographs and the second dataset contained 10,184 micrographs. Both datasets were collected in counting mode and recorded in Electron Event Representation (EER) format using a magnification of 130kx, a pixel size of 0.97 Å, and a nominal dose of 40 e^-^/Å^2^, a defocus range of -0.5 to -2.5 μm, and an energy filter slit width of 10 eV.

The first dataset was processed first in cryoSPARC v4.2 ^56^. Exposures were imported with an EER upsampling factor of 2 and cropped to one-half of their original resolution using Patch Motion Correction followed by CTF correction. Blob picker was first performed to generate 2D classes to use for template picker. 7.9 million particles were picked and extracted with a 270 pixel box size. A subset of particles was used to generate multiple *ab initio* classes. The particle set was cleaned by “decoy classification” through heterogenous refinement, where all 7.9 million particles were assigned to either one of six “junk” classes or one good class (*ab initio* volume resembling a nucleosome). Seven rounds of decoy classification was performed until 441,849 particles remained. The particles were subjected to two rounds of focused 3D classifications followed by two rounds of variability analysis (3DVA) ^57^. The remaining 29,977 particles were subjected to non-uniform refinement (NU-refinement) ^58^, giving a low-resolution structure of USP21 bound to H2AK119ub nucleosome. To improve the resolution of the map, a second dataset was imported as described for the first dataset. TOPAZ was used to train ∼6,000 particles from the map produced from the first dataset to automate particle picking in both datasets, selecting 6.08 million particles. The particles were extracted with a 270 pixel box size and were subjected to four rounds of decoy classification, using one good class and two “junk” classes. Tossed particles subjected to stringent 2D classification to rescue any good particles. Particles that were assigned to classes that were at the Nyquist resolution was reintroduced to the particle stack before a final round of heterogenous refinement, leaving 803,684 particles. The particles were then subjected to two rounds of focused 3D classification with a spherical mask around the USP21 density. This approach originally yielded poor density for USP21 and H2AK119ub, however, after upgrading to cryoSPARC v4.6 which has an improved 3D classification algorithm, we were able to obtain clear density for USP21 and H2AK119ub. After two rounds, the particles were refined by homogenous refinement followed by NU-refinement, yielding a 2.98 Å map from a 83,810 particle stack. A mask was generated around USP21-H2AK119ub and was subjected to local refinement, yielding an USP21-H2AK119ub focus map at a 5.33 Å resolution.

### Model building and refinement

PDB structures of the covalent ubiquitin-USP21 complex (PDB: 3I3T) ^18^ and nucleosome (PDB: 8G6Q) were rigid-body fitted to the map using UCSF ChimeraX ^59^. The model was then rebuilt manually and subjected to real-space refinement using Coot and Phenix^60–62^. The final coordinates were validated using Comprehensive validation (Cryo-EM) in Phenix before deposition in the Protein Data Bank. All figures were generated with UCSF ChimeraX.

### Immunoblotting H2AK119 DUB assay

H2AK119 DUB assays were done at a concentration of 50 nM of USP21 and 400 nM recombinant human H2AK119ub biotinylated 147bp nucleosomes (EpiCypher #16-0395). All reactions were done in 16 uL reactions in USP21 activity assay buffer (25 mM HEPES pH 7.5, 150 mM NaCl, 2 µM ZnCl_2,_ 5 mM DTT). Reactions were done at 30°C for 60 minutes. At each listed timepoint, 4-6 uL of the reaction was quenched in 2x LDS buffer. Quenched reactions were resolved on a 4-12% Bis-Tris Gel (Thermo Fisher) by SDS-PAGE and transferred onto polyvinylidene difluoride (PVDF) membrane using a Trans-Blot Turbo Transfer System (BioRad). Membranes were blocked with 5% milk in 1X Tris-buffered saline with 0.1% Tween-20 (TBS-T) buffer, probed with 1:2500 anti-H2AK119ub (Abcam #ab193203) or 1:2500 anti-H3 (Abcam #ab1791) primary antibodies in 5% milk in 1X TBS-T overnight at 4°C, then probed with 1:5000 anti-rabbit (Invitrogen #31460) horseradish peroxidase (HRP)-conjugated secondary antibody in 5% milk in 1X TBST for 90 minutes at room temperature. Membranes were stained with enhanced chemiluminescence (ECL) reagent and imaged.

For experiments done with phosphorylated USP21, USP21 (2 µM) was first treated with 100 nM of GST-PRKCI (SinoBiological) and GST-MARK (SinoBiological) in kinase activity buffer (25 mM HEPES pH 7.5, 150 mM NaCl, 5 mM MgCl_2_, 20mM ATP, 5 mM DTT). Phosphorylation reactions were done at 30°C for 60 minutes. The phosphorylated protein was then directly diluted in USP21 activity buffer and was used for activity assays as described above.

### K48 di-ubiquitin DUB assay

K48 di-ubiquitin assays were done at a concentration of 100 nM USP21 and 5 µM K48 di-ubiquitin. All reactions were done in USP21 activity assay buffer at 30°C for 60 minutes. At each listed timepoint, 10 uL of the reaction was quenched in 2x LDS buffer. Quenched reactions were resolved on a 4-12% Bis-Tris Gel by SDS-PAGE and stained with Coomassie Blue. For Figure 1C, the band intensity of K48 di-ubiquitin and mono-ubiquitin was measured using Bio-Rad Image Lab and used to calculate fraction of K48 deubiquitinated. The average and standard deviation is reported across technical triplicates.

### Ub-AMC deubiquitination assay

Ub-AMC assays were done with different concentrations of USP21 and 4 µM Ub-AMC in 8 µL reactions. For experiments measuring activity of USP21 truncations (Figure 1B), all reactions were done using 500 nM of USP21. All reactions were performed in USP21 activity assay buffer at 37°C with a Greiner 384-well small volume plate. The AMC fluorescence was measured every 10 seconds using a CLARIOstar Plus Plate Reader. All plots in the main text report the average relative activity and standard deviation across technical triplicates. The normalization to determine relative activity was done using the minimum and maximum mean fluorescence across the triplicates.

Experiments done using the USP21 TEV-cleavable (USP21_I-T-C_) construct, 500 nM USP21_I-T-C_ was incubated with 500 nM TEV protease for 30 minutes at 30°C in USP21 activity assay buffer. Afterwards, the enzyme is then added to Ub-AMC, reaching a final concentration of 250 nM USP21 and 4 µM Ub-AMC. Activity against Ub-AMC was measured as described above.

For experiments done with phosphorylated USP21, USP21 (2 µM) was first treated with 100 nM of GST-PRKCI (SinoBiological) and GST-MARK (SinoBiological) in kinase activity buffer. Phosphorylation reactions were done at 30°C for 60 minutes. The phosphorylated protein was then directly diluted in USP21 activity buffer and was used for Ub-AMC assay, using a concentration of 500 nM USP21 and 4 µM Ub-AMC and measured as described above.

### Electrophoretic mobility shift assays

For EMSAs, 100 nM of 147bp H2AK119ub nucleosome was mixed with either USP21_1-565_ or USP21_196-565_ at 30°C for 60 minutes in EMSA buffer (20 mM HEPES pH 7.5, 50 mM NaCl, 2.5 mM MgCl_2_, 5% Sucrose, 1 mM DTT) in 12 µL reactions. After incubation, 10 uL of the binding reaction was loaded onto a 6% TBE gel at 4°C and ran for 90 minutes at 150 V. The gel was then visualized with SybrGold DNA stain (Thermo).

### Mass spectrometry identification of USP21 phosphorylation sites

USP21 (2 µM) was first treated with 100 nM of GST-PRKCI (SinoBiological) or GST-MARK (SinoBiological) in kinase activity buffer (25 mM HEPES pH 7.5, 150 mM NaCl, 5 mM MgCl_2_, 20mM ATP, 5 mM DTT) at 30°C for 120 minutes. The sample was then flash frozen in liquid nitrogen and stored at -80°C until used for further downstream processing.

Sample preparation and mass spectrometry analysis was carried out by Poochon Scientific (Frederick, Maryland). The sample was divided into two portions and were processed for in-solution trypsin/lysC digestion and chymotrypsin digestion as per SOP-PS-6002 (Standard Operation of Procedure for in-gel Digestion). The digested peptide mixture was then concentrated and desalted using C18 Ziptip as per SOP-PS-6005 (Standard Operation of Procedure for Desalting Digested Peptides). Reconstituted desalted peptides in 20 µl (A) of 0.1% formic acid. 12 µl of peptides was analyzed by 60 min LC/MS/MS run.

The LC/MS/MS analysis of samples were carried out using a Thermo Scientific Orbitrap Q-Exactive Mass Spectrometer and a Thermo Dionex UltiMate 3000 RSLCnano System. Peptide mixture from each sample was loaded onto a peptide trap cartridge at a flow rate of 5 μL/min. The trapped peptides were eluted onto an EasySpray column (Thermo) using a linear gradient of acetonitrile (3-36%) in 0.1% formic acid. The elution duration was 60 min at a flow rate of 0.3 μl/min. Eluted peptides from the EasySpray column were ionized and sprayed into the mass spectrometer, using a Nano-EasySpray Ion Source (Thermo) under the following settings: spray voltage, 1.6 kV, Capillary temperature, 275°C. Raw data file acquired from each sample was searched against human protein sequences database including target USP21 protein sequence using the Proteome Discoverer 2.5 software (Thermo, San Jose, CA) based on the SEQUEST algorithm. Carbamidomethylation (+57.021 Da) of cysteines was fixed modification, Oxidation Met and Q/N-deamidated (+0.98402 Da), and Phospho / +79.966 Da (S, T, Y) were set as dynamic modifications. The minimum peptide length was specified to be five amino acids. The precursor mass tolerance was set to 15 ppm, whereas fragment mass tolerance was set to 0.05 Da. The maximum false peptide discovery rate was specified as 0.01. The resulting Proteome Discoverer Report contains all assembled proteins with peptides sequences and peptide spectrum match counts (PSM#) and MS1 peak area.

### USP21 interaction screen using AlphaPulldown

The sequences of all known interactors of USP21, identified using both low- and high-throughput techniques in the BioGRID database, were extracted and had all duplicates removed. All predictions were performed using the AlphaPulldown package^42^, where all interactors were predicted with both USP21_1-565_ and USP21_196-565_ using AlphaFold2. All AlphaFold2 predictions were performed with three recycles with five models generated for each prediction. The ipTM and PAE scores for the top model of each prediction was used in the analysis.

To identify potential interactors of USP21’s disordered region, we first filtered out all interactors where the ipTM score of the interactor predicted with USP21_1-565_ and USP21_196-565_ was below 0.35, indicating that a low-confidence model. For the remaining interactors, we calculated a ΔipTM score which we define as the difference in ipTM scores of the interactor predicted with USP21_1-565_ and USP21_196-565_. We determined that any interactor with a ΔipTM>0.15 to be a potential candidate interactor of USP21’s IDR.

### H2AK119ub conformation analysis and distance calculations

For the analysis, we extracted the structure of H2AK119ub nucleosome alone^52^, the H2AK119ub-bound structures of SSX1^63^, DNMT3A^64^, PRC2^65^, RYBP-PRC1^66^, UbcH5b-PRC1^67^, PR-DUB^5^, USP16^6^, and USP21 (this study). All H2AK119ub nucleosome containing structures were superimposed onto the crystal structure of nucleosome core particle (PDB-1KX5)^22^ using the core of the histone octamer for alignment. The center of mass using all ubiquitin heavy atoms in each structure was calculated and the distance was measured to the center of mass of all heavy atoms of H2AK119 in 1KX5. All calculations were performed in ChimeraX^59^.

### Structural search of autoinhibited DUBs

All known human ubiquitin specific proteases were visually inspected using the AlphaFold2 database^68^ to identify any DUBs that contain IDRs that overlapped the ubiquitin binding interface. This narrowed the initial list of 54 DUBs down to 18. To ensure that AlphaFold predicts the IDRs to occlude ubiquitin binding, we re-predicted all the DUB structures using AlphaFold3^47^ with and without ubiquitin bound. The apo structure was then analyzed using the Predictomes server^69^. We extracted all DUBs that had contacts between the IDR and catalytic domain which we defined as any interaction that is within 5Å with a PAE cutoff of 20Å and is found in two or more models. To identify DUBs that have IDRs that occlude ubiquitin binding, all DUBs that meet the above criteria were superimposed against the ubiquitin-bound model. Any high-confidence IDR interactions that did not sterically clash with ubiquitin was removed. This narrowed our list of candidate DUBs down to 12.

## Acknowledgements

We thank D. Sousa, D. Ding and K. Cai for their support with cryo-EM sample preparation and data collection at the Beckman Center for Cryo-EM at Johns Hopkins School of Medicine. We thank the Wolberger lab for their insights and discussions on the paper. The authors acknowledge the use of BioRender in the generation of figures present in the manuscript Rahman, S. (2025) https://BioRender.com/87wcwfw. This work was supported by National Institute of General Medical Sciences grants R35GM130393 (C.W.) and National Cancer Institute grants F31CA271743 (S.R.) and F31CA261154 (C.W.H.).

## Author Contributions

S.R. and C.W. conceived and designed the study. S.R., A.G., and M.T.M., prepared all reagents for cryo-EM and deubiquitination assays. S.R. performed all cryo-EM experiments with input from C.W.H. S.R. performed all nucleosome and Ub-AMC activity assays. S.R. and A.G performed di-ubiquitin activity assays. S.R. performed AlphaPulldown screens and structural modeling. C.W. oversaw all aspects of the structure determination, biochemistry, modeling, and data interpretation. S.R. and C.W. wrote the paper, with input from all other authors.

## References

1. Jin, J. et al. The deubiquitinase USP21 maintains the stemness of mouse embryonic stem cells via stabilization of Nanog. Nature Communications 7, 13594 (2016).

2. Nakagawa, T. et al. Deubiquitylation of histone H2A activates transcriptional initiation via trans-histone cross-talk with H3K4 di- and trimethylation. Genes & Development 22, 37–49 (2008).

3. Kulma, M. et al. The ubiquitin-specific protease 21 is critical for cancer cell mitochondrial function and regulates proliferation and migration. Journal of Biological Chemistry 300, 107793 (2024).

4. Jullien, J. et al. Gene Resistance to Transcriptional Reprogramming following Nuclear Transfer Is Directly Mediated by Multiple Chromatin-Repressive Pathways. Molecular Cell 65, 873–884.e8 (2017).

5. Ge, W. et al. Basis of the H2AK119 specificity of the Polycomb repressive deubiquitinase. Nature 616, 176–182 (2023).

6. Ai, H. et al. Structural and mechanistic basis for nucleosomal H2AK119 deubiquitination by single-subunit deubiquitinase USP16. Nature Structural & Molecular Biology 31, 1745–1755 (2024).

7. Wang, H. et al. Role of histone H2A ubiquitination in Polycomb silencing. Nature 431, 873–878 (2004).

8. de Napoles, M. et al. Polycomb Group Proteins Ring1A/B Link Ubiquitylation of Histone H2A to Heritable Gene Silencing and X Inactivation. Developmental Cell 7, 663–676 (2004).

9. Scheuermann, J.C. et al. Histone H2A deubiquitinase activity of the Polycomb repressive complex PR-DUB. Nature 465, 243–247 (2010).

10. Zhu, P. et al. A Histone H2A Deubiquitinase Complex Coordinating Histone Acetylation and H1 Dissociation in Transcriptional Regulation. Molecular Cell 27, 609–621 (2007).

11. Joo, H.-Y. et al. Regulation of cell cycle progression and gene expression by H2A deubiquitination. Nature 449, 1068–1072 (2007).

12. Hong, Y. et al. USP21 Deubiquitinase Regulates AIM2 Inflammasome Activation. The Journal of Immunology 207, 1926–1936 (2021).

13. Liu, J. et al. Ubiquitin-specific protease 21 stabilizes BRCA2 to control DNA repair and tumor growth. Nature Communications 8(2017).

14. Li, W., Cui, K., Prochownik, E.V. & Li, Y. The deubiquitinase USP21 stabilizes MEK2 to promote tumor growth. Cell Death & Disease 9(2018).

15. Xu, G. et al. Ubiquitin-specific Peptidase 21 Inhibits Tumor Necrosis Factor α-induced Nuclear Factor κB Activation via Binding to and Deubiquitinating Receptor-interacting Protein 1. Journal of Biological Chemistry 285, 969–978 (2010).

16. Chen, Y. et al. p38 inhibition provides anti–DNA virus immunity by regulation of USP21 phosphorylation and STING activation. Journal of Experimental Medicine 214, 991–1010 (2017).

17. Ye, Y. et al. Polyubiquitin binding and cross-reactivity in the USP domain deubiquitinase USP21. EMBO reports 12, 350–357 (2011).

18. Ernst, A. et al. A strategy for modulation of enzymes in the ubiquitin system. Science 339, 590–5 (2013).

19. García-Santisteban, I., Bañuelos, S. & Rodríguez, A., Jose. A global survey of CRM1-dependent nuclear export sequences in the human deubiquitinase family. Biochemical Journal 441, 209–217 (2012).

20. Okuda, H. et al. The USP21 Short Variant (USP21SV) Lacking NES, Located Mostly in the Nucleus In Vivo, Activates Transcription by Deubiquitylating ubH2A In Vitro. PLoS ONE 8, e79813 (2013).

21. Morgan, M.T. et al. Structural basis for histone H2B deubiquitination by the SAGA DUB module. Science 351, 725–728 (2016).

22. Davey, C.A., Sargent, D.F., Luger, K., Maeder, A.W. & Richmond, T.J. Solvent mediated interactions in the structure of the nucleosome core particle at 1.9 Å resolution. Journal of Molecular Biology 319, 1097–1113 (2002).

23. Thomas, J.F. et al. Structural basis of histone H2A lysine 119 deubiquitination by Polycomb repressive deubiquitinase BAP1/ASXL1. Science Advances 9(2023).

24. McGinty, R.K. & Tan, S. Principles of nucleosome recognition by chromatin factors and enzymes. Curr Opin Struct Biol 71, 16–26 (2021).

25. Skrajna, A. et al. Comprehensive nucleosome interactome screen establishes fundamental principles of nucleosome binding. Nucleic Acids Res 48, 9415–9432 (2020).

26. Zhou, X. et al. Exploring genomic alteration in pediatric cancer using ProteinPaint. Nature Genetics 48, 4–6 (2016).

27. Sondka, Z. et al. COSMIC: a curated database of somatic variants and clinical data for cancer. Nucleic Acids Research 52, D1210–D1217 (2024).

28. Gersch, M. et al. Distinct USP25 and USP28 Oligomerization States Regulate Deubiquitinating Activity. Mol Cell 74, 436–451 e7 (2019).

29. Sauer, F. et al. Differential Oligomerization of the Deubiquitinases USP25 and USP28 Regulates Their Activities. Mol Cell 74, 421–435 e10 (2019).

30. Zeqiraj, E. et al. Higher-Order Assembly of BRCC36-KIAA0157 Is Required for DUB Activity and Biological Function. Mol Cell 59, 970–83 (2015).

31. Deng, M. et al. Deubiquitination and Activation of AMPK by USP10. Mol Cell 61, 614–624 (2016).

32. Huang, X. et al. Deubiquitinase USP37 is activated by CDK2 to antagonize APC(CDH1) and promote S phase entry. Mol Cell 42, 511–23 (2011).

33. Samara, N.L. et al. Structural insights into the assembly and function of the SAGA deubiquitinating module. Science 328, 1025–9 (2010).

34. Cohn, M.A. et al. A UAF1-containing multisubunit protein complex regulates the Fanconi anemia pathway. Mol Cell 28, 786–97 (2007).

35. Heride, C. et al. The centrosomal Deubiquitylase USP21 regulates Gli1 transcriptional activity and stability. Journal of Cell Science, jcs.188516 (2016).

36. Urbé, S. et al. Systematic survey of deubiquitinase localization identifies USP21 as a regulator of centrosome- and microtubule-associated functions. Molecular Biology of the Cell 23, 1095–1103 (2012).

37. Sowa, M.E., Bennett, E.J., Gygi, S.P. & Harper, J.W. Defining the Human Deubiquitinating Enzyme Interaction Landscape. Cell 138, 389–403 (2009).

38. Boldt, K. et al. An organelle-specific protein landscape identifies novel diseases and molecular mechanisms. Nature Communications 7, 11491 (2016).

39. Luck, K. et al. A reference map of the human binary protein interactome. Nature 580, 402–408 (2020).

40. Oughtred, R. et al. The BioGRID database: A comprehensive biomedical resource of curated protein, genetic, and chemical interactions. Protein Science 30, 187–200 (2021).

41. Evans, R. et al. Protein complex prediction with AlphaFold-Multimer. (Cold Spring Harbor Laboratory, 2021).

42. Yu, D., Chojnowski, G., Rosenthal, M. & Kosinski, J. AlphaPulldown-a python package for protein-protein interaction screens using AlphaFold-Multimer. Bioinformatics 39(2023).

43. Morrison, D.K. The 14-3-3 proteins: integrators of diverse signaling cues that impact cell fate and cancer development. Trends Cell Biol 19, 16–23 (2009).

44. Madeira, F. et al. 14-3-3-Pred: improved methods to predict 14-3-3-binding phosphopeptides. Bioinformatics 31, 2276–2283 (2015).

45. Johnson, J.L. et al. An atlas of substrate specificities for the human serine/threonine kinome. Nature 613, 759–766 (2023).

46. Johnson, J.L. et al. An atlas of substrate specificities for the human serine/threonine kinome. Nature 613, 759–766 (2023).

47. Abramson, J. et al. Accurate structure prediction of biomolecular interactions with AlphaFold 3. Nature 630, 493–500 (2024).

48. Komander, D. Mechanism, specificity and structure of the deubiquitinases. Subcell Biochem 54, 69–87 (2010).

49. Morgan, M.T. et al. Structural basis for histone H2B deubiquitination by the SAGA DUB module. Science 351, 725–8 (2016).

50. Luger, K., Rechsteiner, T.J. & Richmond, T.J. Preparation of nucleosome core particle from recombinant histones. Methods Enzymol 304, 3–19 (1999).

51. Nodelman, I.M., Patel, A., Levendosky, R.F. & Bowman, G.D. Reconstitution and Purification of Nucleosomes with Recombinant Histones and Purified DNA. Curr Protoc Mol Biol 133, e130 (2020).

52. Hicks, W., Chad, et al. Ubiquitinated histone H2B as gatekeeper of the nucleosome acidic patch. Nucleic Acids Research 52, 9978–9995 (2024).

53. Lowary, P.T. & Widom, J. New DNA sequence rules for high affinity binding to histone octamer and sequence-directed nucleosome positioning. J Mol Biol 276, 19–42 (1998).

54. Makde, R.D., England, J.R., Yennawar, H.P. & Tan, S. Structure of RCC1 chromatin factor bound to the nucleosome core particle. Nature 467, 562–6 (2010).

55. Dyer, P.N. et al. Reconstitution of nucleosome core particles from recombinant histones and DNA. Methods Enzymol 375, 23–44 (2004).

56. Punjani, A., Rubinstein, J.L., Fleet, D.J. & Brubaker, M.A. cryoSPARC: algorithms for rapid unsupervised cryo-EM structure determination. Nat Methods 14, 290–296 (2017).

57. Punjani, A. & Fleet, D.J. 3D variability analysis: Resolving continuous flexibility and discrete heterogeneity from single particle cryo-EM. J Struct Biol 213, 107702 (2021).

58. Punjani, A., Zhang, H. & Fleet, D.J. Non-uniform refinement: adaptive regularization improves single-particle cryo-EM reconstruction. Nat Methods 17, 1214–1221 (2020).

59. Pettersen, E.F. et al. UCSF ChimeraX: Structure visualization for researchers, educators, and developers. Protein Sci 30, 70–82 (2021).

60. Emsley, P. & Cowtan, K. Coot: model-building tools for molecular graphics. Acta Crystallogr D Biol Crystallogr 60, 2126–32 (2004).

61. Adams, P.D. et al. PHENIX: a comprehensive Python-based system for macromolecular structure solution. Acta Crystallogr D Biol Crystallogr 66, 213–21 (2010).

62. Chen, V.B. et al. MolProbity: all-atom structure validation for macromolecular crystallography. Acta Crystallogr D Biol Crystallogr 66, 12–21 (2010).

63. Tong, Z. et al. Synovial sarcoma X breakpoint 1 protein uses a cryptic groove to selectively recognize H2AK119Ub nucleosomes. Nat Struct Mol Biol 31, 300–310 (2024).

64. Chen, X. et al. Structural basis for the H2AK119ub1-specific DNMT3A-nucleosome interaction. Nat Commun 15, 6217 (2024).

65. Kasinath, V. et al. JARID2 and AEBP2 regulate PRC2 in the presence of H2AK119ub1 and other histone modifications. Science 371, eabc3393 (2021).

66. Ciapponi, M., Karlukova, E., Schkolziger, S., Benda, C. & Muller, J. Structural basis of the histone ubiquitination read-write mechanism of RYBP-PRC1. Nat Struct Mol Biol 31, 1023–1027 (2024).

67. Ai, H. et al. Synthetic E2-Ub-nucleosome conjugates for studying nucleosome ubiquitination. Chem 9, 1221–1240 (2023).

68. Tunyasuvunakool, K. et al. Highly accurate protein structure prediction for the human proteome. Nature 596, 590–596 (2021).

69. Schmid, E.W. & Walter, J.C. Predictomes: A classifier-curated database of AlphaFold-modeled protein-protein interactions. bioRxiv (2024).

