## Supplemental Figures for "Mechanism of USP21 autoinhibition and histone H2AK119 deubiquitination"

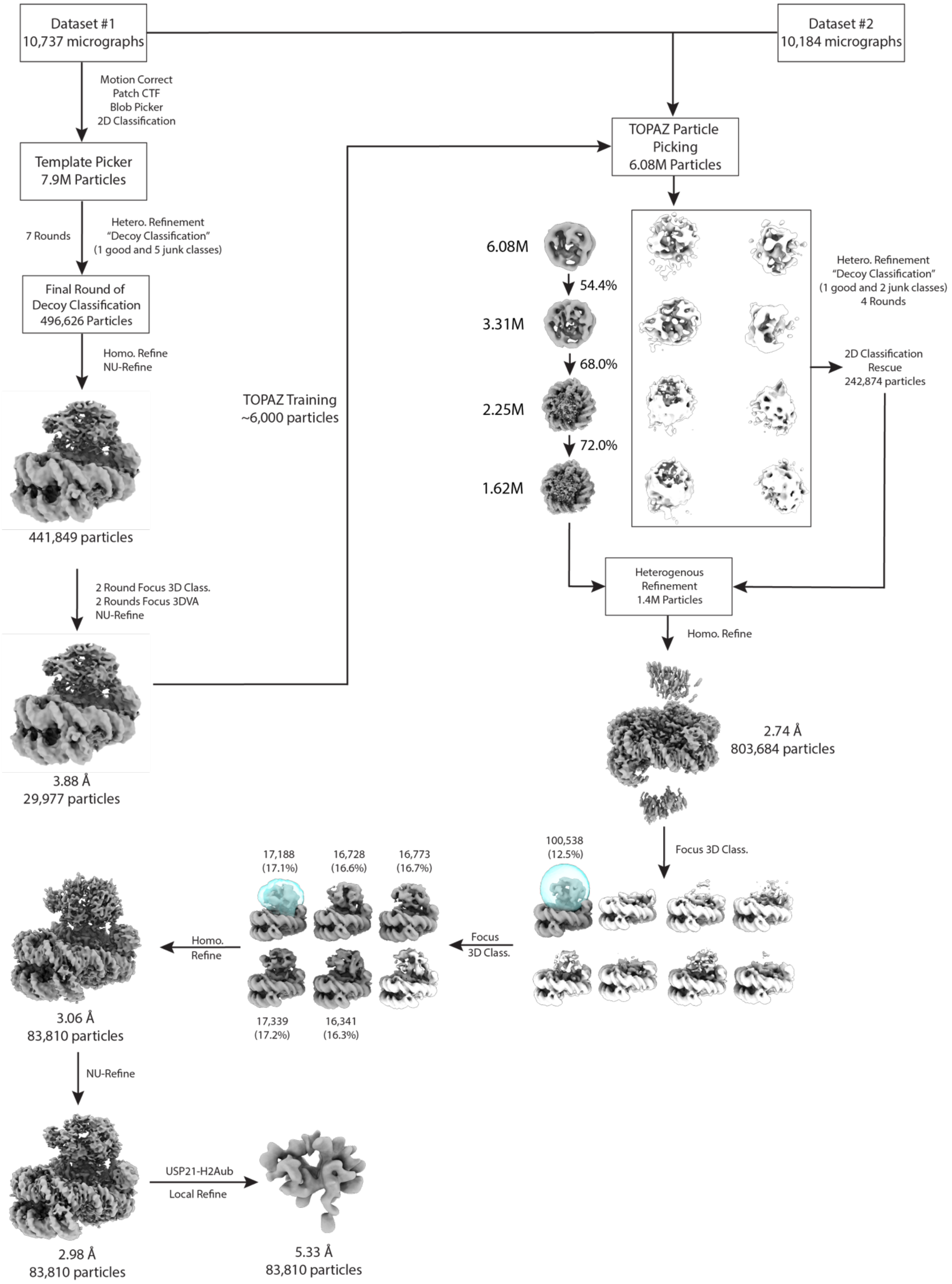


**Supplemental Figure 1- Data Processing of USP21**

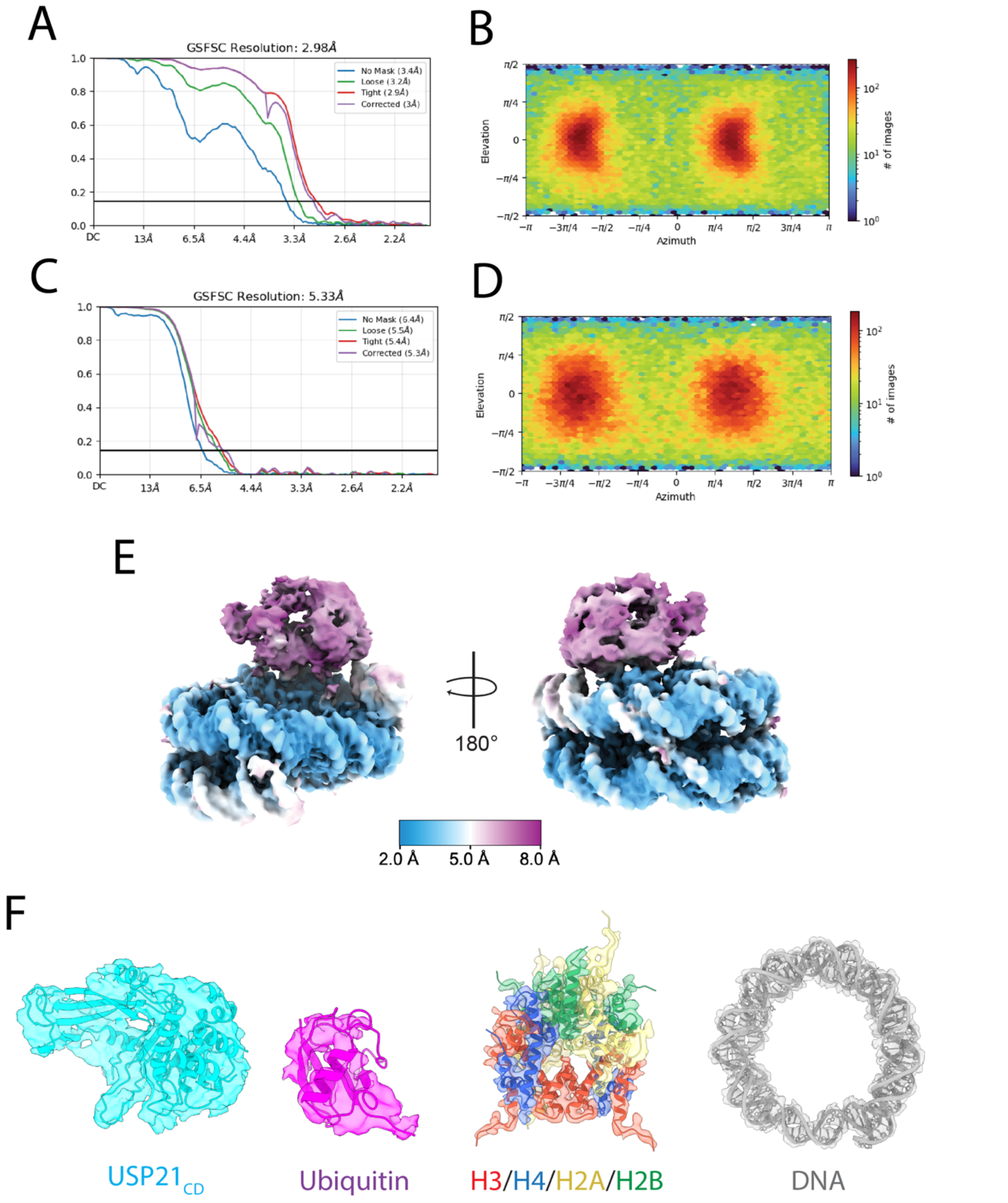


**Supplemental Figure 2- Map Validation and Resolution Calculations** (A) Fourier Shell Correlation (FSC) curve and (B) particle distribution orientation (B) of the USP21-H2AK119ub nucleosome map. (C) Fourier Shell Correlation (FSC) curve and (D) particle distribution orientation of the USP21-focused map. (E) Local resolution of USP21-H2AK119ub nucleosome map. (F) Model-map fit of USP21, ubiquitin, histone octamer, and nucleosome DNA.

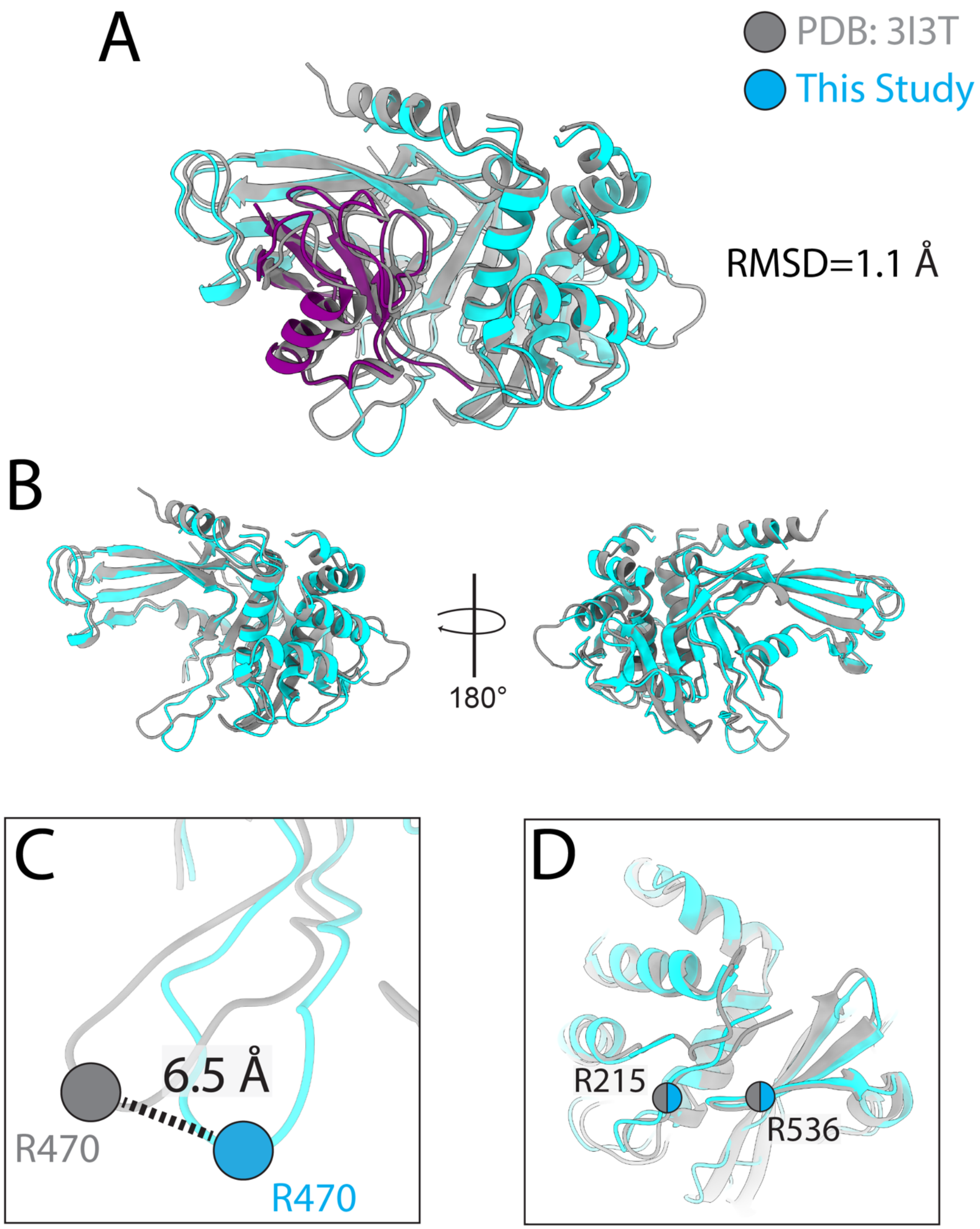


**Supplemental Figure 3- Comparison of USP21-Ub interaction from crystallography and EM.** (A) Superimposition of USP21-Ub interaction from the crystal structure (PDB: 3I3T) onto the USP21-Ub model in this study. (B) Comparison of USP21 structure in the crystal structure and cryo-EM model. Comparison of USP21 (C) acidic patch and (D) DNA-binding residues in the crystal structure and cryo-EM model.

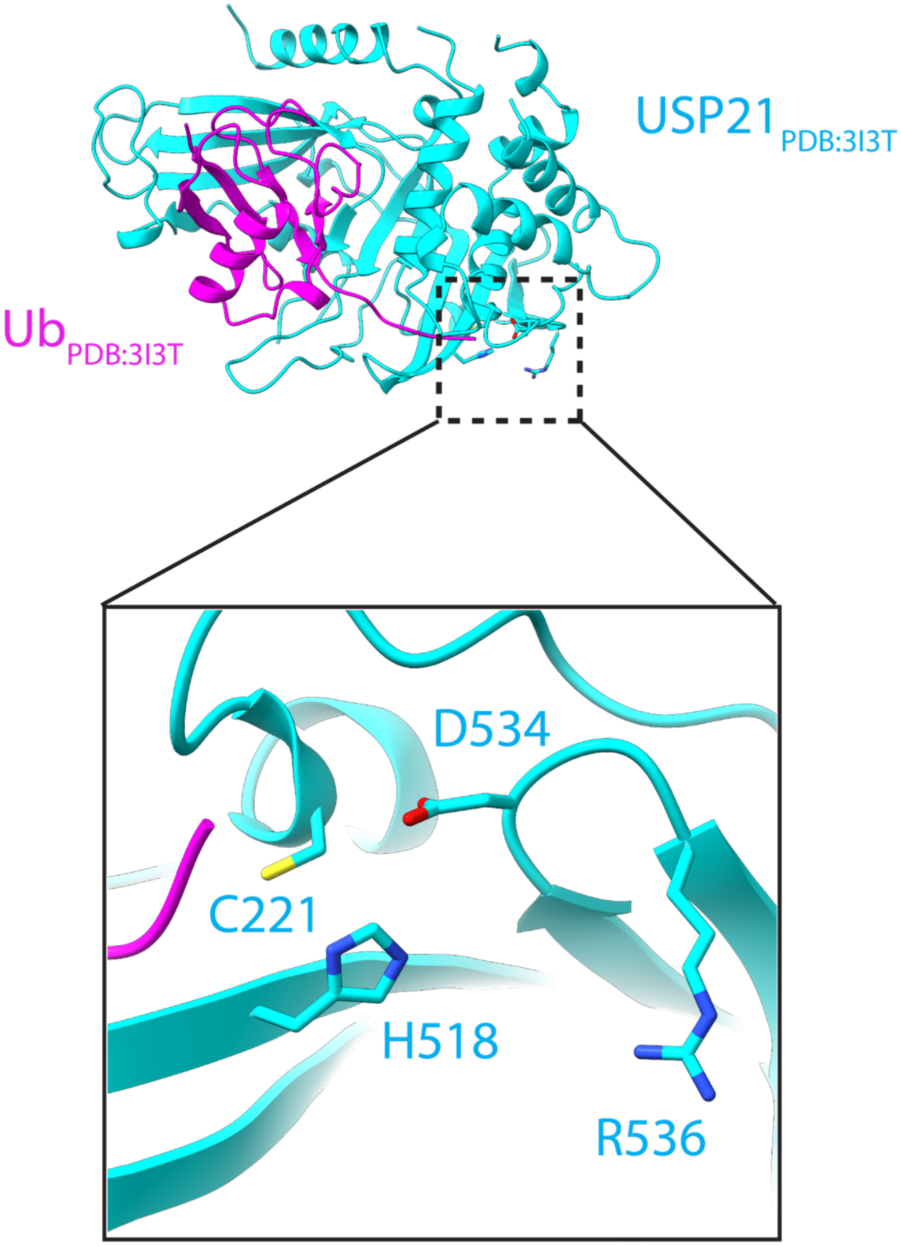


**Supplemental Figure 4. USP21$_{R536}$ is positioned near USP21's catalytic triad**. Zoom-in on the catalytic site of USP21 (PDB- 3I3T) showing the positioning of R536 in respect to the catalytic cysteine (C221) and triad residues H518 and D534.

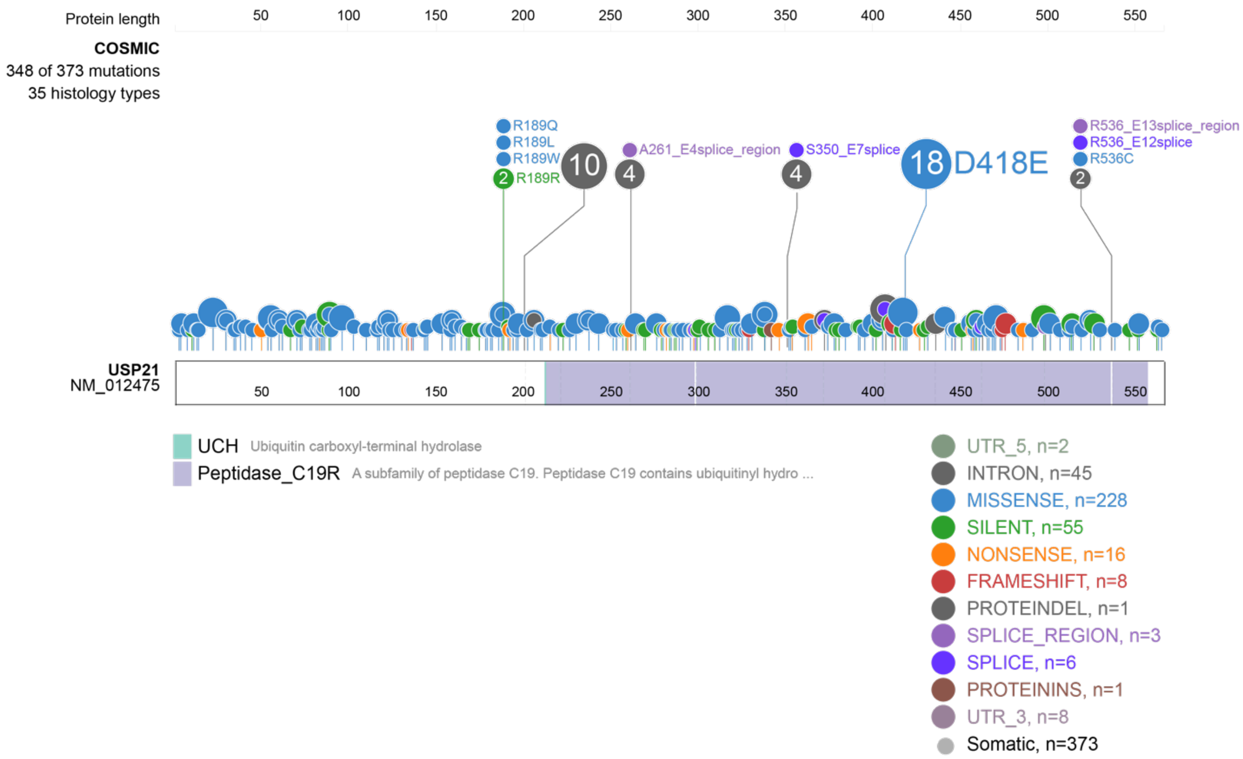


**Supplemental Figure 5. Structural analysis of USP21 cancer mutations.** Structural mapping of all known USP21 mutations identified from the COSMIC database using St. Jude's ProteinPaint server[26].

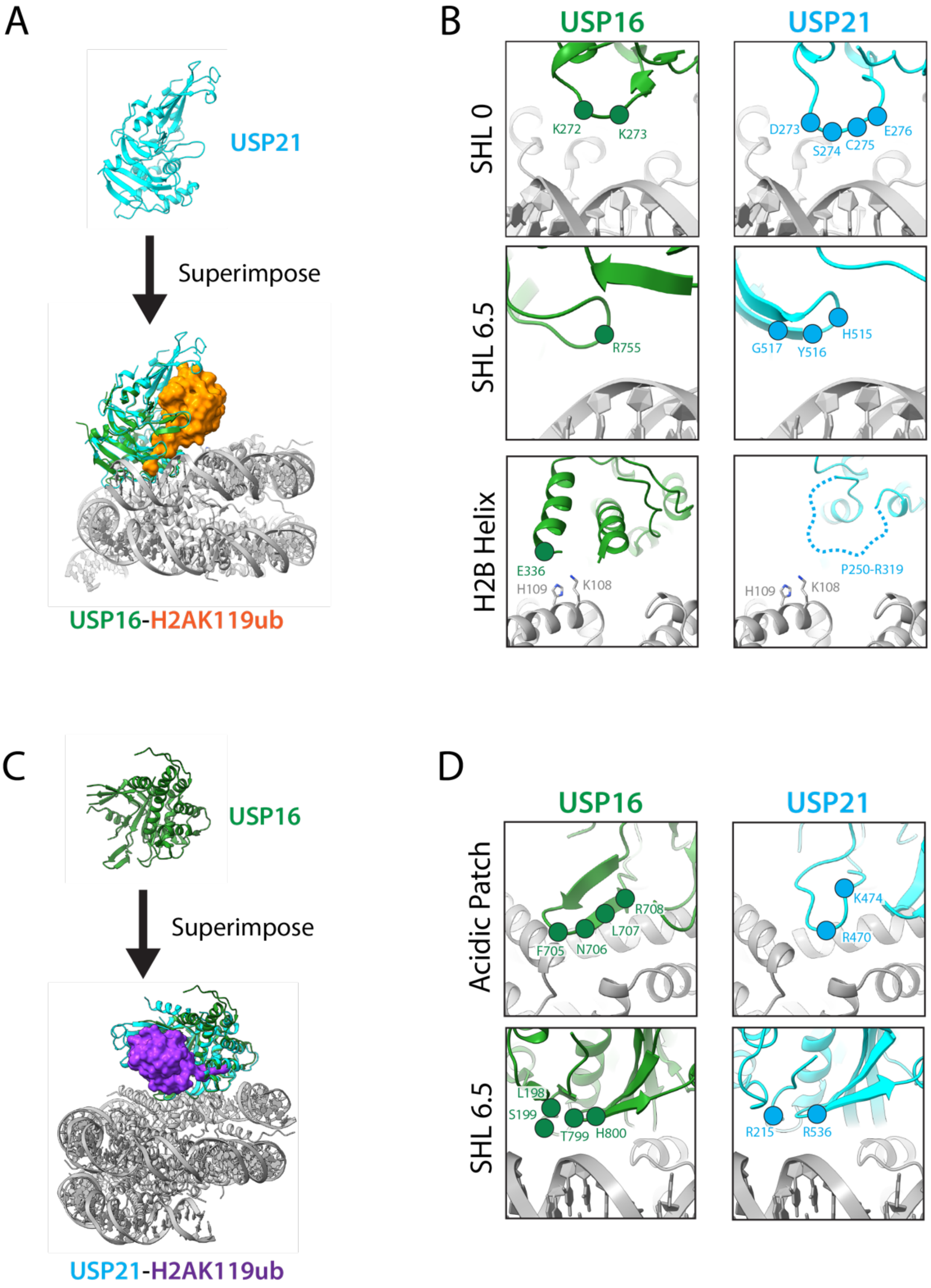


**Supplemental Figure 6. Comparison of USP16 and USP21 H2AK119ub nucleosome engagement**. (A) Combability of USP21 binding to H2AK119ub through USP16's mode of binding. (B) Zoom-in on structure alignment in (A). Comparison of USP16 and USP21's interactions with H2AK119ub nucleosome, critical for USP16's mode of binding. (C) Combability of USP16 binding to H2AK119ub through USP21's mode of binding. (B) Zoom-in on structure alignment in (C). Comparison of USP16 and USP21's interactions with H2AK119ub nucleosome, critical for USP21's mode of binding.

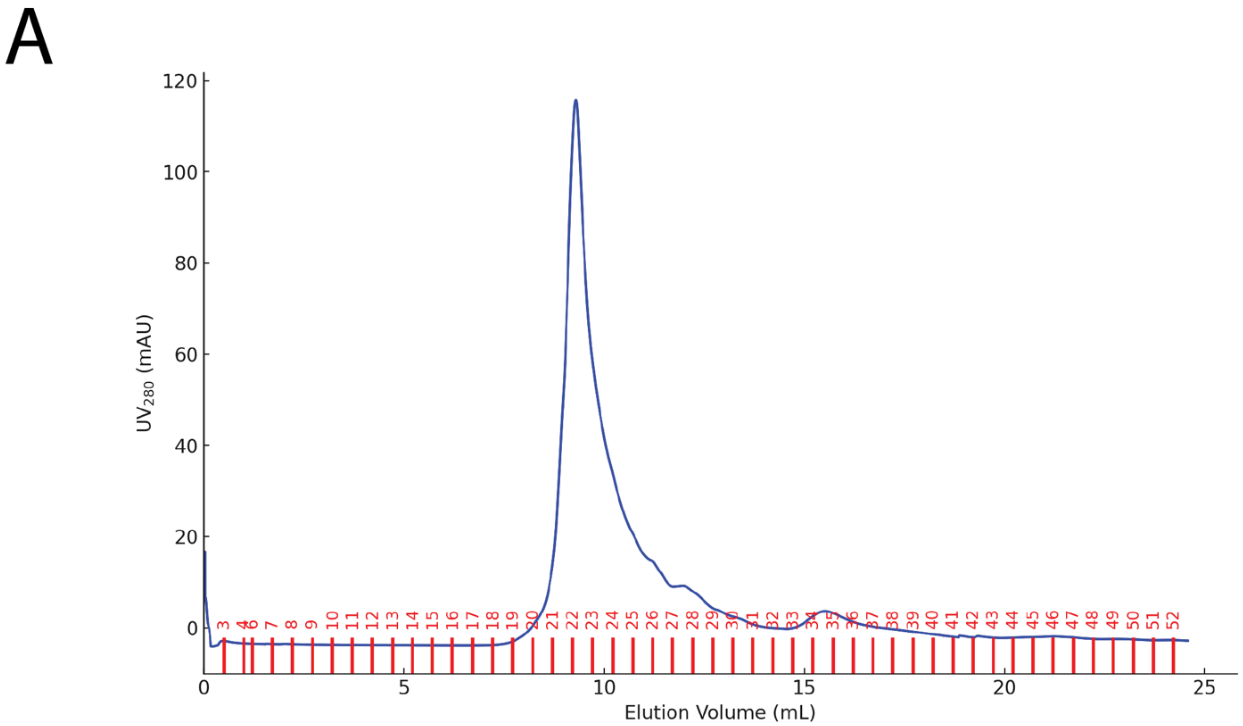


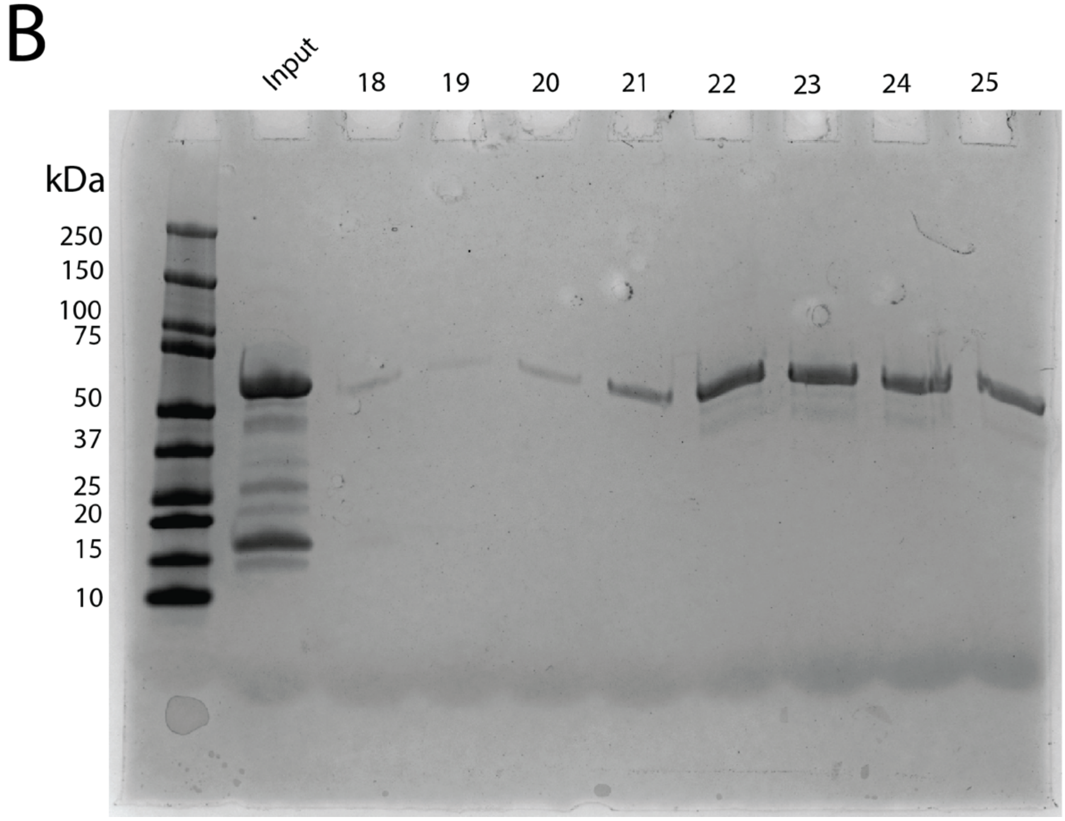


**Supplemental Figure 7. Purity and olgiomerization state of USP21 over size-exclusion chromatography (SEC).** (A) SEC trace of full-length USP21 on a Superdex 75 10/300 column. (B) Coomassie blue analysis of USP21 protein after SEC purification.

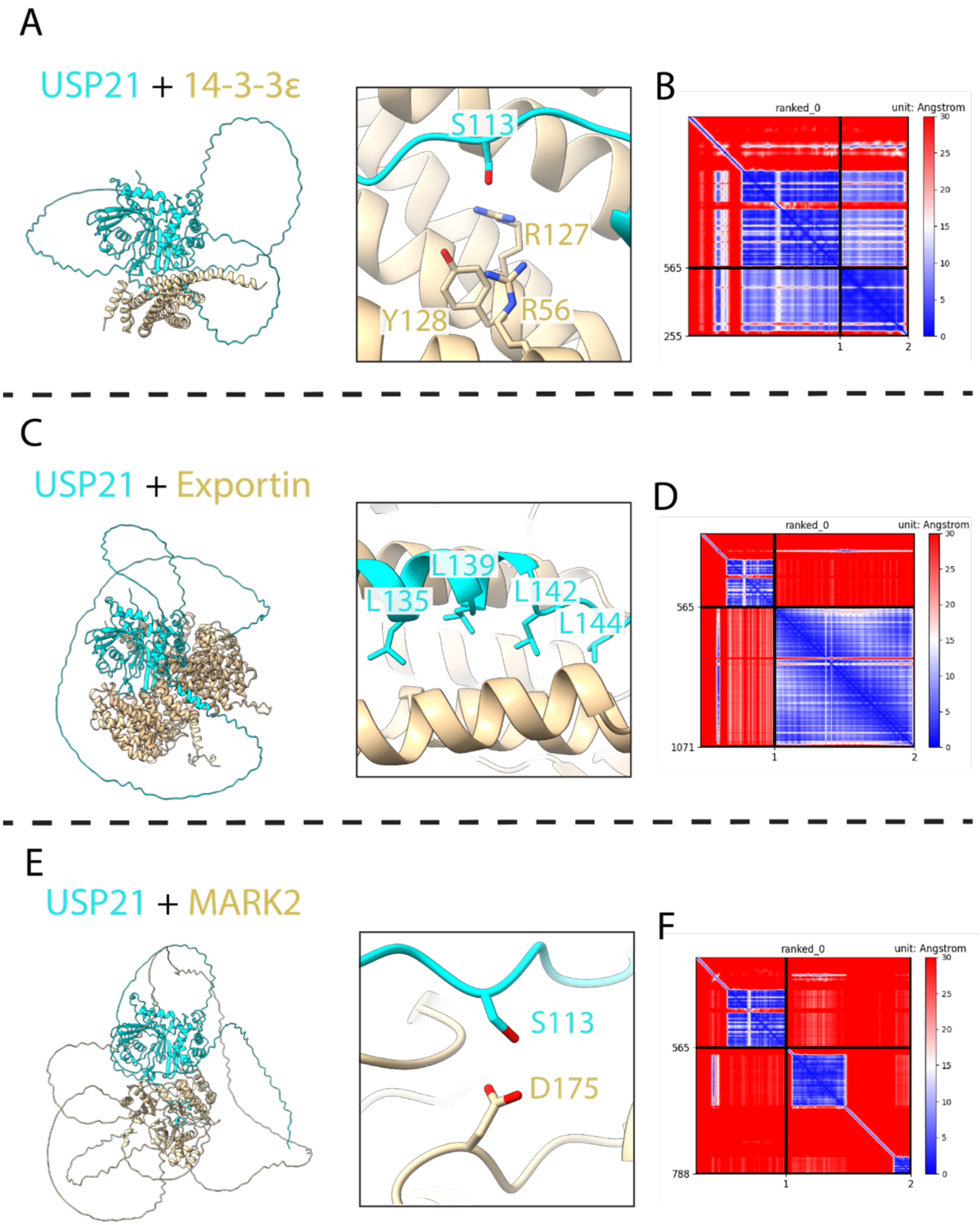


**Supplemental Figure 8. Additional high-confidence interactors of USP21's IDR predicted by AlphaFold-Multimer (AF-M).** (A) AF-M prediction of USP21 and 14-3-3ε with zoom-in on the phosphoserine binding pocket of 14-3-3ε, recognizing $USP21_{S113}$. (B) Predicted Aligned Error (PAE) of the top AF-M model of the USP21 and 14-3-3ε complex. (C) AF-M prediction of USP21 and Exportin with zoom-in on USP21's nuclear export signal sequence (LXXXLXXLXL). (D) PAE of the top AF-M model of the USP21 and Exportin complex. (E) AF-M prediction of USP21 and MARK2 with zoom-in on MARK2's active site, poised to phosphorylated $USP21_{S113}$. (F) PAE of the top AF-M model of the USP21 and MARK2 complex.

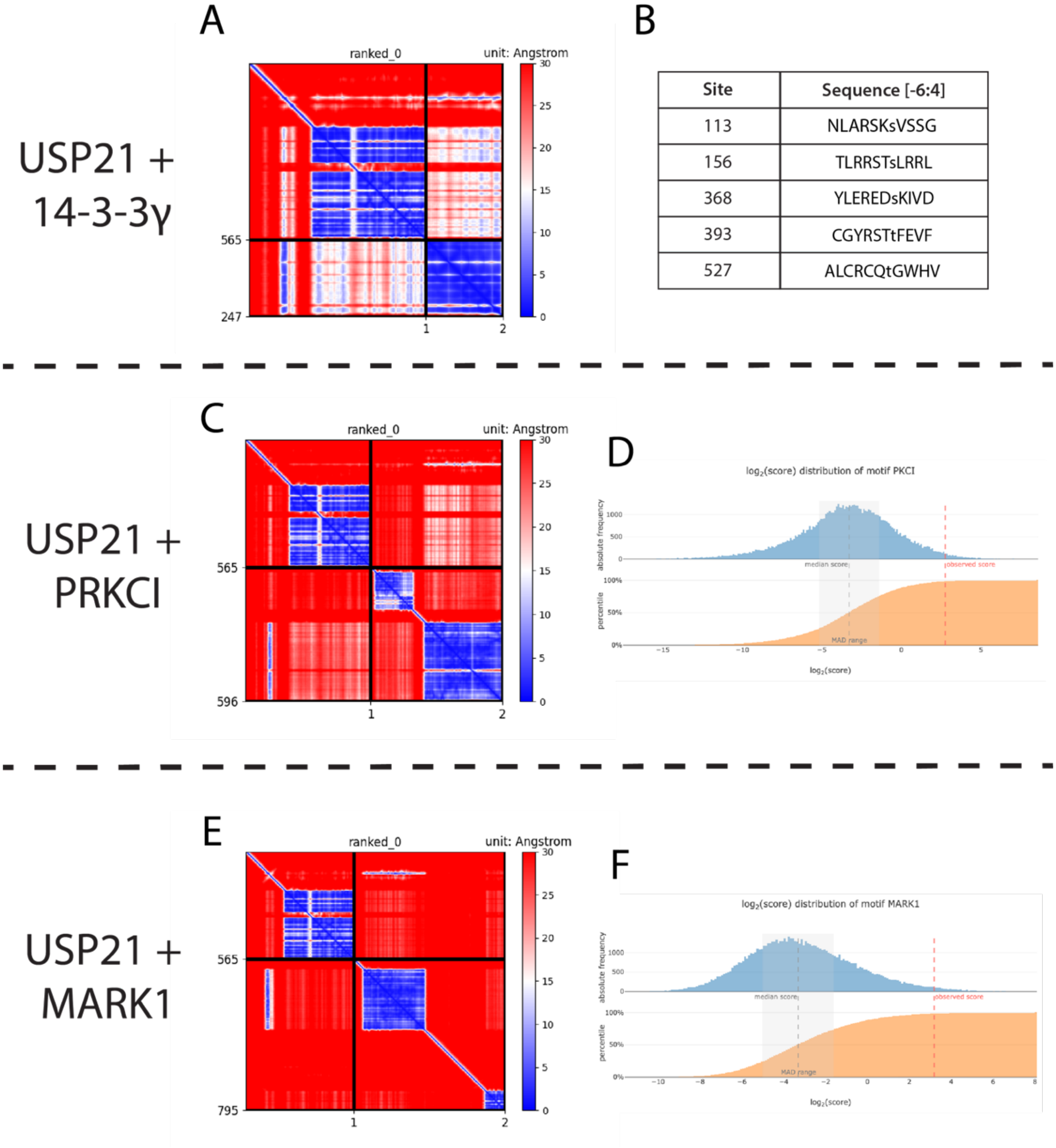


**Supplemental Figure 9. AlphaFold-Multimer (AF-M) model confidence and sequence scoring for 14-3-3 binding and kinase phosphorylation.** (A) Predicted Aligned Error (PAE) of the top AF-M model of the USP21 and 14-3-3γ complex. (B) List of predicted high-confidence 14-3-3 binding sites on USP21, with the key phosphorylated residue in lowercase using the 14-3-3 Pred server[44]. (C) PAE of the top AF-M model of the USP21 and PRKCI complex. (D) $Log_2$ score of $USP21_{S113}$ for phosphorylation by PRKCI (PKCI) predicted by Kinase Library. (E) PAE of the top AF-M model of the USP21 and MARK1 complex. (F) $Log_2$ score of $USP21_{S115}$ for phosphorylation by MARK1 predicted by Kinase Library.

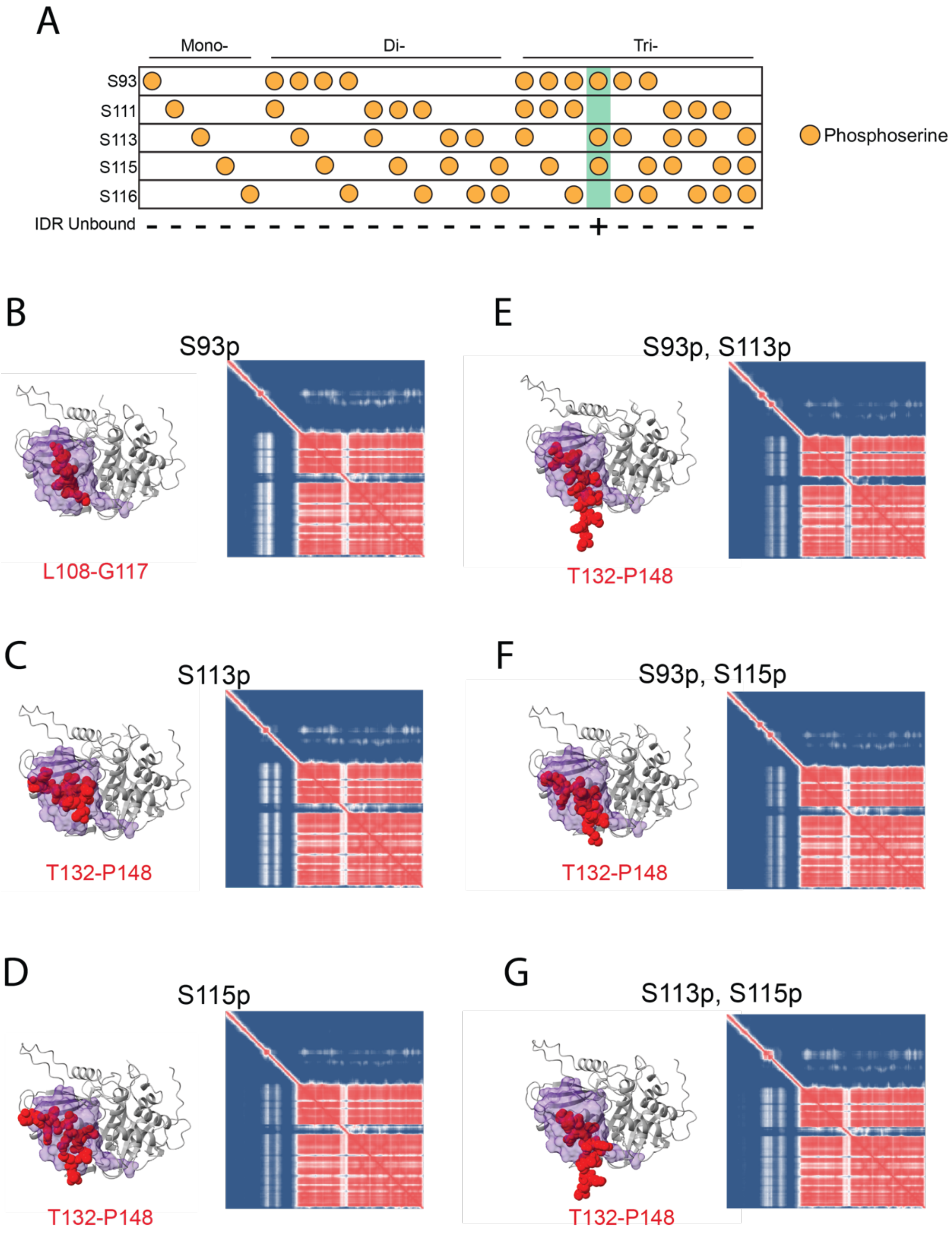


**Supplemental Figure 10. AlphaFold3 (AF3) modeling of USP21 in various phosphorylation states.** (A) Summary of all phosphorylated forms of USP21 predicted by AF3 and their impact on repositioning $USP21_{IDR}$ away from the ubiquitin-binding interface. (B-G) AF3 model and Predicted Aligned Error of USP21 phosphorylated at S93, S113, and/or S115 (apo). Residues of $USP21_{IDR}$ that obscure ubiquitin-binding are shown in red. All models are aligned to the ubiquitin-bound AF3 model of USP21 to show ubiquitin binding orientation (purple).

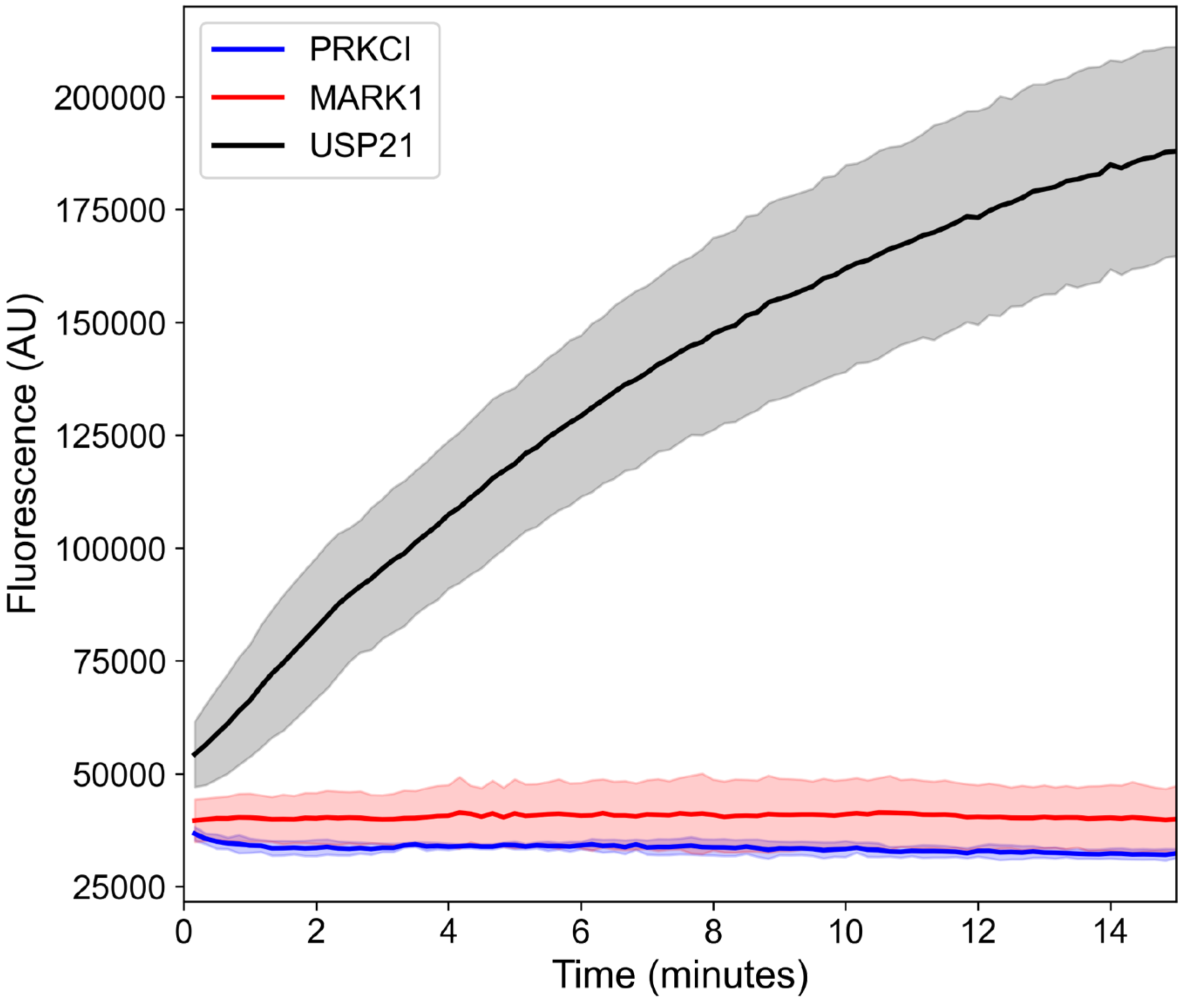


**Supplemental Figure 11. Activity of PRKCI and MARK1 against ubiquitin-aminomethylcoumarin (Ub-AMC).** Time-course curve of Ub-AMC cleavage by PRKCI and MARK1, showing neither kinase have activity against Ub-AMC or contain any contaminating deubiquitinases. Ub-AMC activity is shown in comparison to full-length USP21.

A

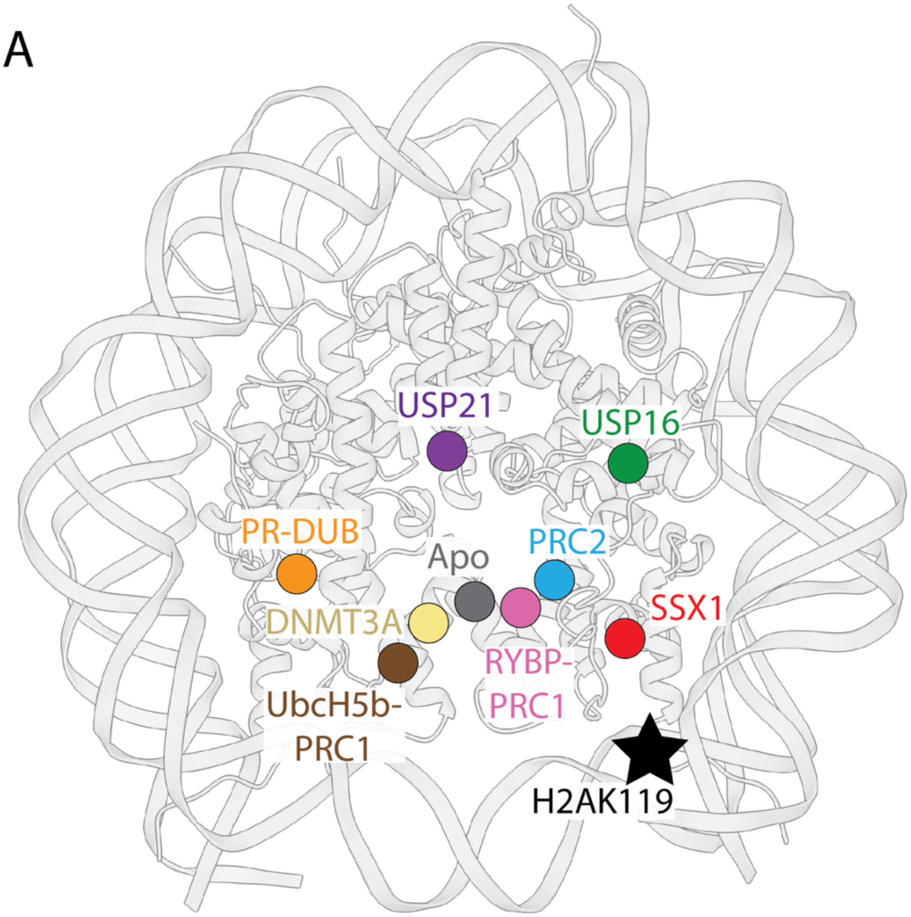


B

| Structure | $Distance_{Ub-H2AK119}$ (Å) |
| --- | --- |
| SSX1 | 21.6 |
| Apo | 25.5 |
| DNMT3A | 27.5 |
| PRC2 | 29.3 |
| RYBP-PRC1 | 33.1 |
| UbcH5b-PRC1 | 34.5 |
| PR-DUB | 43.2 |
| USP16 | 44.1 |
| USP21 | 46.2 |

**Supplementary Figure 12. Conformational diversity of H2AK119ub positioning.** (A) Structural mapping of ubiquitin positioning across all H2AK119ub-containing structures. Each point represents the center of mass of ubiquitin in each structure. (B) Distance calculation between the center of mass of ubiquitin and H2AK119 across all H2AK119ub-containing structures, in angstroms.

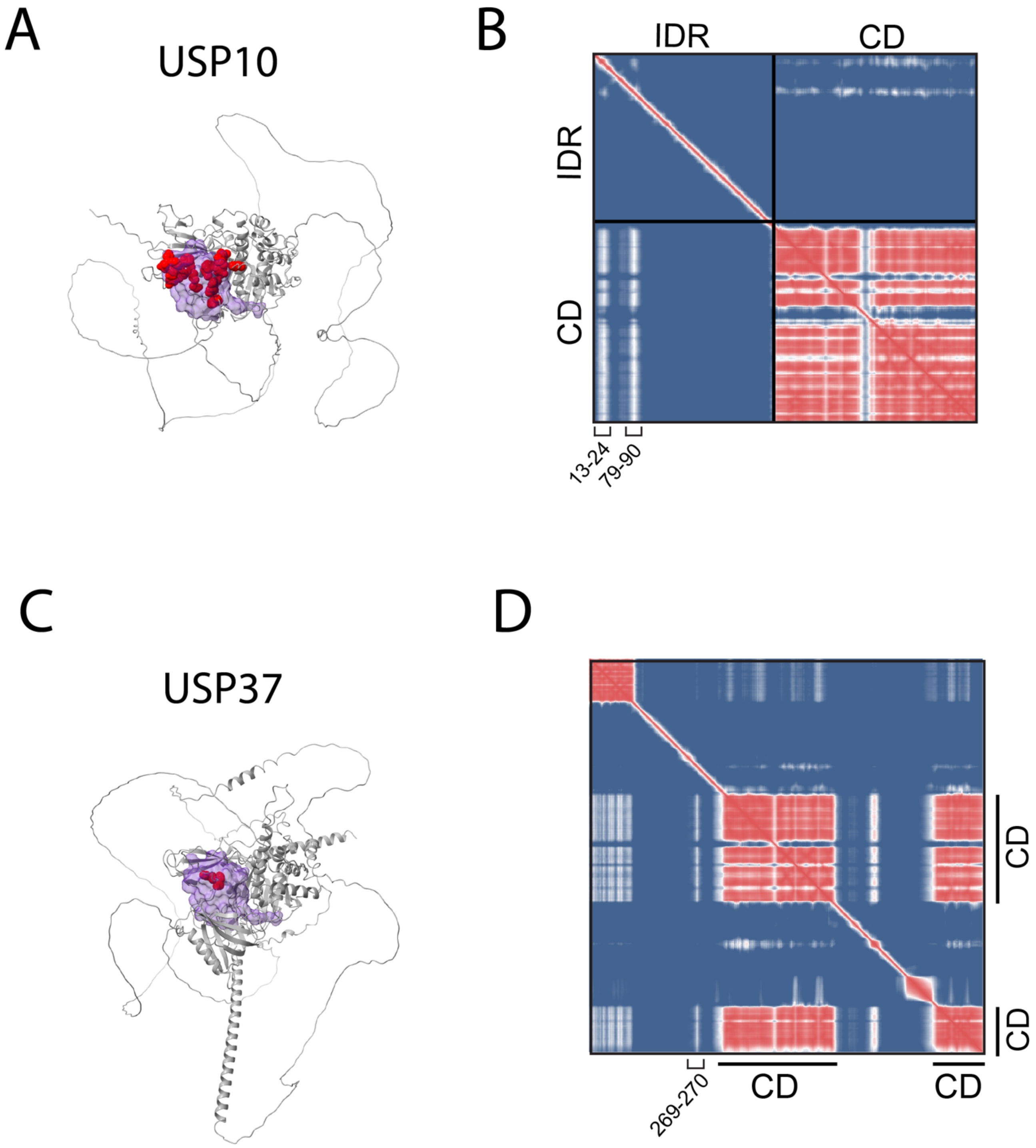


**Supplemental Figure 13. AlphaFold3 (AF3) modeling of USP10 and USP37 phosphorylation.** (A) AF3 model of USP10 with autoinhibitory residues shown as red spheres and shown in respect to the ubiquitin-bound model predicted by AF3 (purple). (B) Predicted Aligned Error (PAE) of the top AF3 model of USP37. (C) AF3 model of USP10 with autoinhibitory residues shown as red spheres and shown in respect to the ubiquitin-bound model predicted by AF3 (purple). (D) Predicted Aligned Error (PAE) of the top AF3 model of USP10. All five models of USP10 and USP37 gave similar structure predictions and PAE scores.

**List of supplemental files (.xls):**

***Supplementary Table 1- Structure and map validation scores***

***Supplementary Table 2- List of USP21 mutation from COSMIC database***

***Supplementary Table 3- USP21 AlphaPulldown Results***

***Supplementary Table 4- Mass spectrometry analysis of USP21 phosphorylation sites***

***Supplementary Table 5- Predictome analysis of all ubiquitin specific proteases***
